# Post-infection immune response in adults with COVID-19 with and without nirmatrelvir-ritonavir treatment and virologic rebound

**DOI:** 10.64898/2026.06.22.733737

**Authors:** Dibya Ghimire, Chloe M. Hasund, John Bianchi Colangeli, May Y. Liew, Emory Abar, Mamadou Barry, Andrew Alexandrescu, Benjamin L. Kotzen, Zahra Reynolds, Eliza Passell, Chase Mandell, Zoie Smith-Sparks, Karry Su, Julie Boucau, Owen T. Glover, Brooke Leeman, Gregory E. Edelstein, Manish C. Choudhary, Yijia Li, Naomi Patel, Jeffrey A. Sparks, Amy K. Barczak, Jonathan Z. Li, Mark J. Siedner, Jacob E. Lemieux

**Affiliations:** Massachusetts General Hospital, Boston, Massachusetts; Department of Immunology and Infectious Diseases, Harvard T.H. Chan School of Public Health, Boston, MA, USA; Infectious Disease and Microbiome Program, Broad Institute, Cambridge, MA, USA; Broad Institute, Cambridge, Massachusetts; Ragon Institute of Mass General Brigham, MIT, and Harvard, Cambridge, Massachusetts; Brigham and Women’s Hospital, Boston, Massachusetts; University of Pittsburgh Medical Center, Pittsburgh, Pennsylvania; Harvard Medical School, Boston, Massachusetts

## Abstract

**Background:** Nirmatrelvir-ritonavir (N-R) reduces morbidity and mortality from COVID-19 in high-risk individuals; however, N-R use has been associated with risk of SARS-CoV-2 virologic rebound. The mechanisms contributing to virologic rebound after N-R treatment are currently unknown. One plausible mechanism is that antiviral treatment may alter the development of immune responses to SARS-CoV-2 infection, thereby contributing to rebound after cessation of therapy.

**Methods:** We profiled immune responses in a case-ascertained, longitudinal, prospective cohort of ambulatory individuals with COVID-19. Participants were grouped according to whether virological rebound occurred and whether they had received N-R treatment. We assessed antibody, T-cell, and innate responses.

**Results:** We observed no differences in the binding or neutralizing antibody, T-cell, and innate immune responses between participants with virologic rebound compared to participants without virologic rebound. N-R use was associated with slightly weaker antibody responses overall, even after adjustment for immunosuppression.

**Conclusion:** Virologic rebound after N-R treatment is likely driven by non-immune mechanisms.

**Funding:** The project was supported by the National Institutes of Health (R01 AI 138801) and the Massachusetts Consortium on Pathogen Readiness. Its contents are solely the responsibility of the authors and do not necessarily represent the official views of the NIH.

## Introduction

Nirmatrelvir-ritonavir (N-R) is approved by the FDA to treat adults (18 years and above) diagnosed with mild to moderate COVID-19 at high risk for progression to severe disease. The drug is also approved under an Emergency Use Authorization (EUA) for the treatment of adults and pediatric patients (12 years of age and older weighing at least 40 kg) with mild to moderate COVID-19 and at high risk for progression to hospitalization or death(1). Nirmatrelvir works by inhibiting SARS-CoV-2’s main protease (Mpro)(2–4), which cleaves the polyproteins Pp1 and Pp1ab into components of the SARS-CoV-2 replication-transcription complex. Ritonavir acts as a pharmacological enhancer by inhibiting cytochrome P450 3A, prolonging the half-life of nirmatrelvir. In initial studies, N-R significantly reduced the risk of hospitalization and death rate by 70-90%(5–11), although more recent evaluations suggest that the beneficial effects are likely more modest, possibly due to the current landscape of population immunity and circulating variants.(12, 13)

N-R treatment is associated with increased incidence of virologic rebound(14–17). Virologic rebound occurs in 15-30% of N-R treated subjects treated with the standard 5-day regimen. Longer duration regimens of 10 and 15 days reduce the rate of rebound(18), and retreatment upon virologic rebound is also safe and effective(19).

The mechanisms by which N-R treatment can result in virologic rebound are not currently known. Several hypotheses are plausible, including pharmacologic blunting of the immune response needed for complete viral clearance, emergence of viral mutations that lead to N-R resistance, and preservation of target cells that enables resumption of replication when drug pressure is removed. Among these hypotheses, acquired N-R resistance appears unlikely to explain rebound because independent groups have found an absence of M^pro^-resistance mutations among subjects with rebound(14, 16, 20). Mathematical modeling demonstrates that preservation of target cells, either by a robust innate immune response or initiation of N-R near the time of symptom onset, coupled with incomplete viral clearance, can account for rebound dynamics(21). Consistent with this hypothesis, treatment of SARS-CoV-2-infected cells *in vitro* with N-R and another M^Pro^ protease inhibitor, ensitrelvir, results in an infectious virus that persists longer than treatment with remdesivir, a polymerase inhibitor(22). However, neither is a proven mechanism of rebound in humans with COVID-19. The possibility of N-R treatment blunting the immune response to infection has been suggested by one study(23), but another has found no effect(24); however, both studies are small (n= 3 in Panza et al.(23), n=6 in Epling et al.(24)), and larger studies are needed.

In the parent POSt-VaccInaTIon Viral CharactEristics Study (POSITIVES) prospective cohort, 20.8% of participants who received N-R treatment experienced virologic rebound, whereas only 1.8% of untreated participants experienced rebound(14). The present study represents an immunologic sub-analysis of this cohort and includes participants with available longitudinal blood specimens, as well as additional eligible participants enrolled after publication of the original cohort report. To investigate the immunologic mechanisms potentially associated with virologic rebound following N-R treatment, we comprehensively evaluated both humoral and cellular immune responses during acute and post-acute SARS-CoV-2 infection. We quantified the abundance of antiviral antibodies directed against Spike and Nucleocapsid proteins, measured neutralizing antibody titers against multiple SARS-CoV-2 variants, and analyzed the antibody kinetics over time. To assess cell-mediated immunity, we performed ex-vivo fluorospot assays to quantify IFN-ɣ producing T-cells following Spike peptide pool stimulation. In parallel, to further characterize innate immune activation and inflammatory signaling, we profiled cytokine and chemokine responses using multiplex Luminex assays. By integrating these complementary immune profiling approaches across clinically stratified cohorts, we attempted to determine whether virologic rebound after N-R treatment is associated with impaired antiviral immunity, delayed immune resolution, or persistent inflammatory activation. Collectively, our findings suggest that virologic rebound following N-R treatment is not strongly associated with major defects in systemic humoral, cellular, or innate immune responses.

## RESULTS

### Cohort characteristics

We investigated the immune response among 153 participants with acute COVID-19 enrolled in the POSITIVES study(14, 15, 20, 25–30). Of these participants, 75 were treated with N-R and 76 were untreated. The demographics and key viral characteristics of the cohort are summarized in Table 1. The N-R treated group was similar to the non-treated group in terms of age, sex, race, ethnicity, and vaccination profiles but contained a higher proportion of immunosuppressed participants. The rate of virologic rebound in the N-R-treated participants with available samples was (13/75, 17.3%).

**Table 1:**
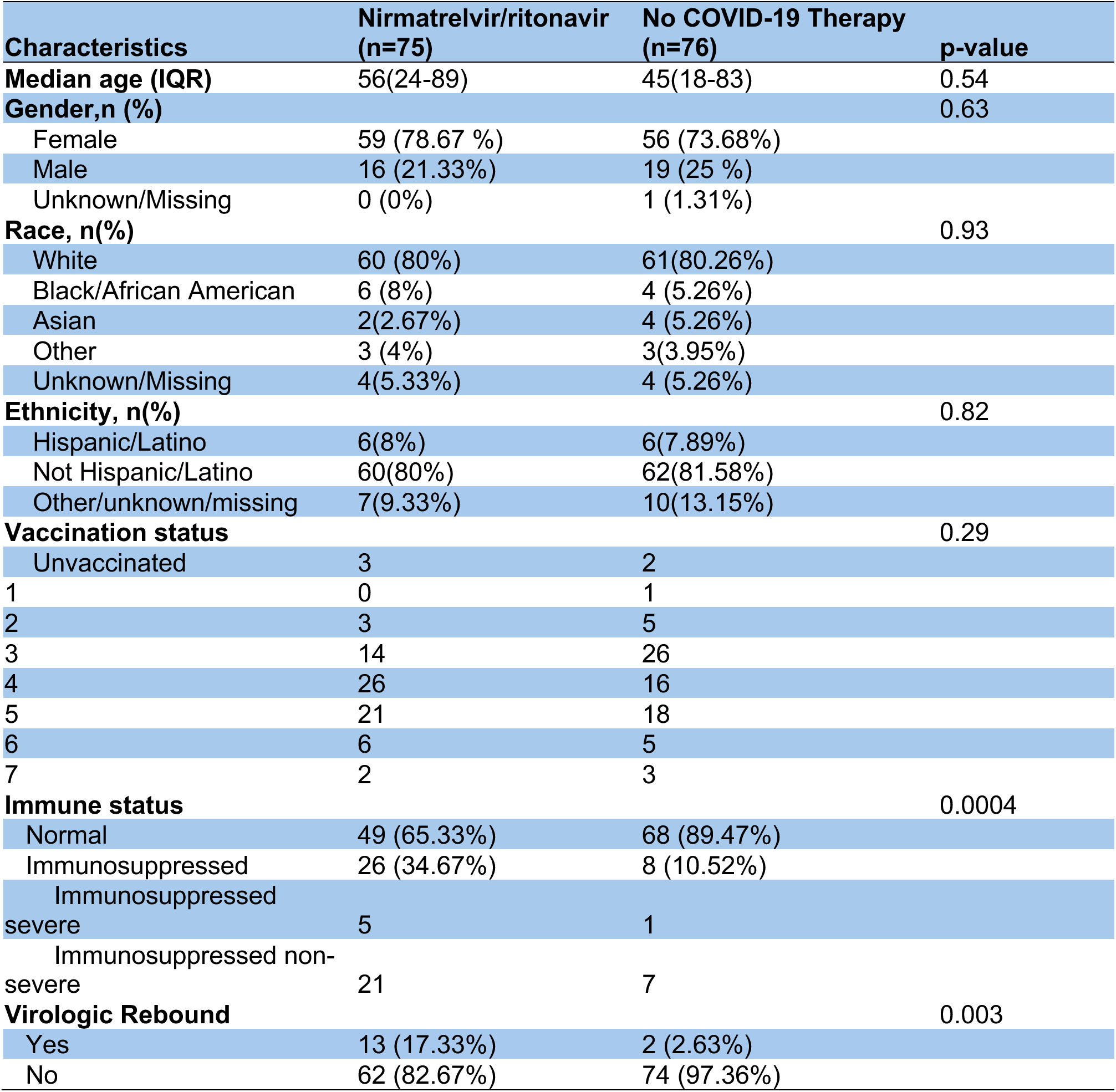
Characteristics of the subjects in the cohort.

| Characteristics | Nirmatrelvir/ritonavir<br>(n=75) | No COVID-19 Therapy<br>(n=76) | p-value |
| --- | --- | --- | --- |
| **Median age (IQR)** | 56(24-89) | 45(18-83) | 0.54 |
| **Gender, n (%)** |  |  | 0.63 |
| Female | 59 (78.67 %) | 56 (73.68%) |  |
| Male | 16 (21.33%) | 19 (25 %) |  |
| Unknown/Missing | 0 (0%) | 1 (1.31%) |  |
| **Race, n(%)** |  |  | 0.93 |
| White | 60 (80%) | 61(80.26%) |  |
| Black/African American | 6 (8%) | 4 (5.26%) |  |
| Asian | 2(2.67%) | 4 (5.26%) |  |
| Other | 3 (4%) | 3(3.95%) |  |
| Unknown/Missing | 4(5.33%) | 4 (5.26%) |  |
| **Ethnicity, n(%)** |  |  | 0.82 |
| Hispanic/Latino | 6(8%) | 6(7.89%) |  |
| Not Hispanic/Latino | 60(80%) | 62(81.58%) |  |
| Other/unknown/missing | 7(9.33%) | 10(13.15%) |  |
| **Vaccination status** |  |  | 0.29 |
| Unvaccinated | 3 | 2 |  |
| 1 | 0 | 1 |  |
| 2 | 3 | 5 |  |
| 3 | 14 | 26 |  |
| 4 | 26 | 16 |  |
| 5 | 21 | 18 |  |
| 6 | 6 | 5 |  |
| 7 | 2 | 3 |  |
| **Immune status** |  |  | 0.0004 |
| Normal | 49 (65.33%) | 68 (89.47%) |  |
| Immunosuppressed | 26 (34.67%) | 8 (10.52%) |  |
| Immunosuppressed severe | 5 | 1 |  |
| Immunosuppressed non-severe | 21 | 7 |  |
| **Virologic Rebound** |  |  | 0.003 |
| Yes | 13 (17.33%) | 2 (2.63%) |  |
| No | 62 (82.67%) | 74 (97.36%) |  |

### Antibody responses to SARS-CoV-2 are similar in subjects with and without virologic rebound

Anti-SARS-CoV-2 Nucleocapsid antibody titers were low during Timepoint 1 (acute infection) and increased nearly 15-fold at Timepoint 2 (post-acute infection phase) in both participants who did and did not experience virologic rebound (Figure 1A), as well as N-R treated participants and those who received no COVID-19 therapy (Figure 1B). Titers subsequently declined at Timepoints 3 and 4. Anti-SARS-CoV-2 Nucleocapsid antibody levels in the N-R-treated group were consistently lower than the untreated group at all four timepoints; however, the differences were not statistically significant except for Timepoint 3 (Figure 1B). When anti-SARS-CoV-2 Nucleocapsid antibody titers were stratified jointly by COVID-19 therapy and virologic rebound status, similar antibody kinetics were observed across all the groups from Timepoint 1 to Timepoint 2 (Figure 1C).

**Figure 1.**
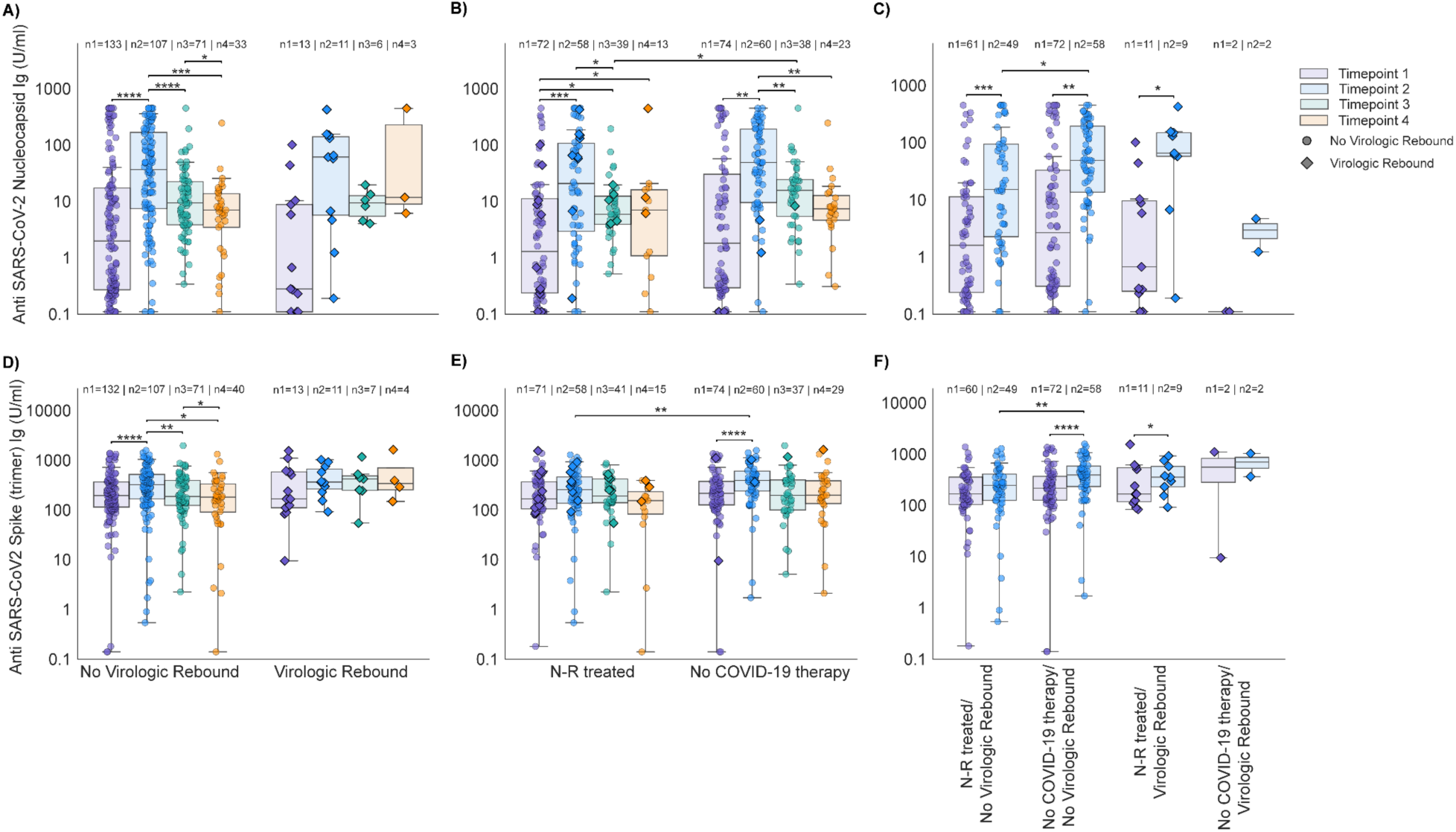
Nirmatrelvir-ritonavir treatment and virologic rebound shape anti–SARS-CoV-2 antibody responses over time. Anti–SARS-CoV-2 Nucleocapsid **(A–C)** and Anti–SARS-CoV-2 Spike Trimer **(D–F)** antibody concentrations (U/mL) measured at four timepoints after COVID-19 detection. **(A, D)** All participants stratified by virologic rebound status (no virologic rebound vs. virologic rebound). **(B, E)** Participants stratified by treatment: N-R treated vs. no COVID-19 therapy. **(C, F)** Participants stratified jointly by treatment and virologic rebound status, yielding four groups: no virologic rebound / N-R treated, no virologic rebound / no COVID-19 therapy, virologic rebound / N-R treated, and virologic rebound / no COVID-19 therapy. Timepoint 1 represents the acute phase (0–15 days after detection), Timepoint 2 the post-acute phase (15–60 days), Timepoint 3 approximately 6 months, and Timepoint 4 approximately 1 year after COVID-19 detection. Sample sizes per timepoint (n1–n4) are indicated above each group. Multiple comparisons across timepoints within a group were performed using the Wilcoxon matched-pairs signed rank test with Benjamini-Hochberg false discovery rate (FDR) correction. Comparisons between independent groups were performed using the Mann-Whitney U test. Box plots show the 25th, 50th, and 75th percentiles; whiskers indicate maximum and minimum values. All data points are shown; y-axes are on a logarithmic scale. Diamond symbols represent virologic rebounders. Symbols above the brackets indicate the degree of significance. No asterisks is nonsignificant,**** p<0.0001, *** *P* < 0.001, ** *P* < 0.01 and * *P* < 0.05.

A similar trajectory was observed in anti-SARS-CoV-2 Spike trimer antibody titer (Figure 1D); however, anti-Spike antibody levels at Timepoint 1 were elevated, likely reflecting prior vaccination, consistent with the well-documented persistence of vaccine-induced humoral immunity for months to years following COVID-19 vaccination (31, 32). Anti-SARS-CoV-2 Spike trimer antibody responses were strong in both N-R-treated and untreated groups at all four timepoints. Although the untreated group showed visibly higher titers than the N-R-treated group across all timepoints, this difference reached statistical significance only at Timepoint 2 (Figure 1E). Joint stratification of the cohort by treatment and virologic rebound status revealed a significant difference in Spike antibody titers at Timepoint 2 between the individuals with no virologic rebound but different treatment status, similar to the pattern observed for anti-Nucleocapsid titers (Figure 1 F).

Immunosuppressed participants were more likely than immunocompetent participants to receive N-R, likely due to higher risk for severe clinical outcomes. To evaluate immunosuppression as a possible confounding variable in the analysis, we compared antibody levels by immune status. Both immunocompetent individuals and immunosuppressed individuals showed expected normal longitudinal kinetics (Figure S2 A and C). No significant differences were observed between two groups at any timepoints; there was a trend toward weaker anti-Nucleocapsid responses at Timepoint 2 (Figure S2 C) among immunosuppressed participants. When immunosuppressed individuals were further stratified by immunosuppression severity as defined by detailed medical record review (27), those with severe immunosuppression demonstrated impaired humoral immunity compared to immunocompetent and non-severe immunosuppressed individuals, characterized by lower anti-Spike and anti-Nucleocapsid antibody titres and altered longitudinal antibody kinetics (Figure S2B, D) compared to immunocompetent individuals.

In addition to unadjusted group comparisons, we performed multivariable linear regression to evaluate the independent effects of demographic and clinical covariates on humoral immune responses (Table 2). Antibody levels were significantly higher at approximately Timepoint 2 (15-60 days post-infection) compared to Timepoint 1 (baseline) (p <0.001). Consistent with the trends observed in unadjusted analysis, time since infection was the primary determinant of humoral immune responses. No significant associations were observed with age, sex, vaccination status, immunosuppression, or N-R treatment among participants who did not experience virologic rebound.

**Table 2:** Regression coefficients for the fitted linear mixed effects model for anti-SARS-CoV-2 nucleocapsid antibody titers.

| Predictors | Estimates (B) | SE | *p-value* |
| --- | --- | --- | --- |
| (Intercept) | 0.65 | 0.36 | 0.07 |
| Dummy timepoint [1] | 0.83 | 0.18 | *** |
| Participant type [Received N-R, no virologic rebound] | -0.47 | 0.25 | 0.07 |
| Sex [Male] | 0.04 | 0.23 | 0.86 |
| Immunosuppressed [Yes] | -0.19 | 0.25 | 0.45 |
| Age category [<40 years] | 0.09 | 0.26 | 0.73 |
| Age category [>=60 years] | -0.12 | 0.25 | 0.64 |
| Vaccinations | 0.03 | 0.07 | 0.65 |
| Dummy timepoint [1] x Participant type [Received N-R, no virologic rebound] | 0.20 | 0.26 | 0.45 |

Reference groups: Dummy time point 0 (baseline), did not receive N-R and did not have viral rebound, female, not immunosuppressed, age category (40-59)
Vaccinations were included in the model as a number, and ranged from 0 to 7.

### Variant-specific neutralization is similar among subjects with and without virologic rebound and N-R treatment is associated with slightly reduced neutralization

We next assessed the neutralizing capacity of anti-SARS-CoV-2 antibodies in serum against different SARS-CoV-2 variants (WT and 5 different variants B.1.1.7 (Alpha), B.1.351 (Beta), P.1 (Gamma), B.1.617.2 (Delta) and B.1.1.529 (Omicron)) using bead-based Luminex assays across timepoints 1 and 2. Given the time period of the study, all the individuals studied were infected with SARS-CoV-2 Omicron variant, and the overall neutralizing capacity of their serum was lower against Omicron compared to other tested variants. Stratification by COVID-19 treatment showed significant differences in treatment groups; however, virologic rebound status did not significantly alter the serum neutralizing capacity against Omicron variant or other tested variants (Figure 2 A,B, Supplementary data part 1).

**Figure 2:**
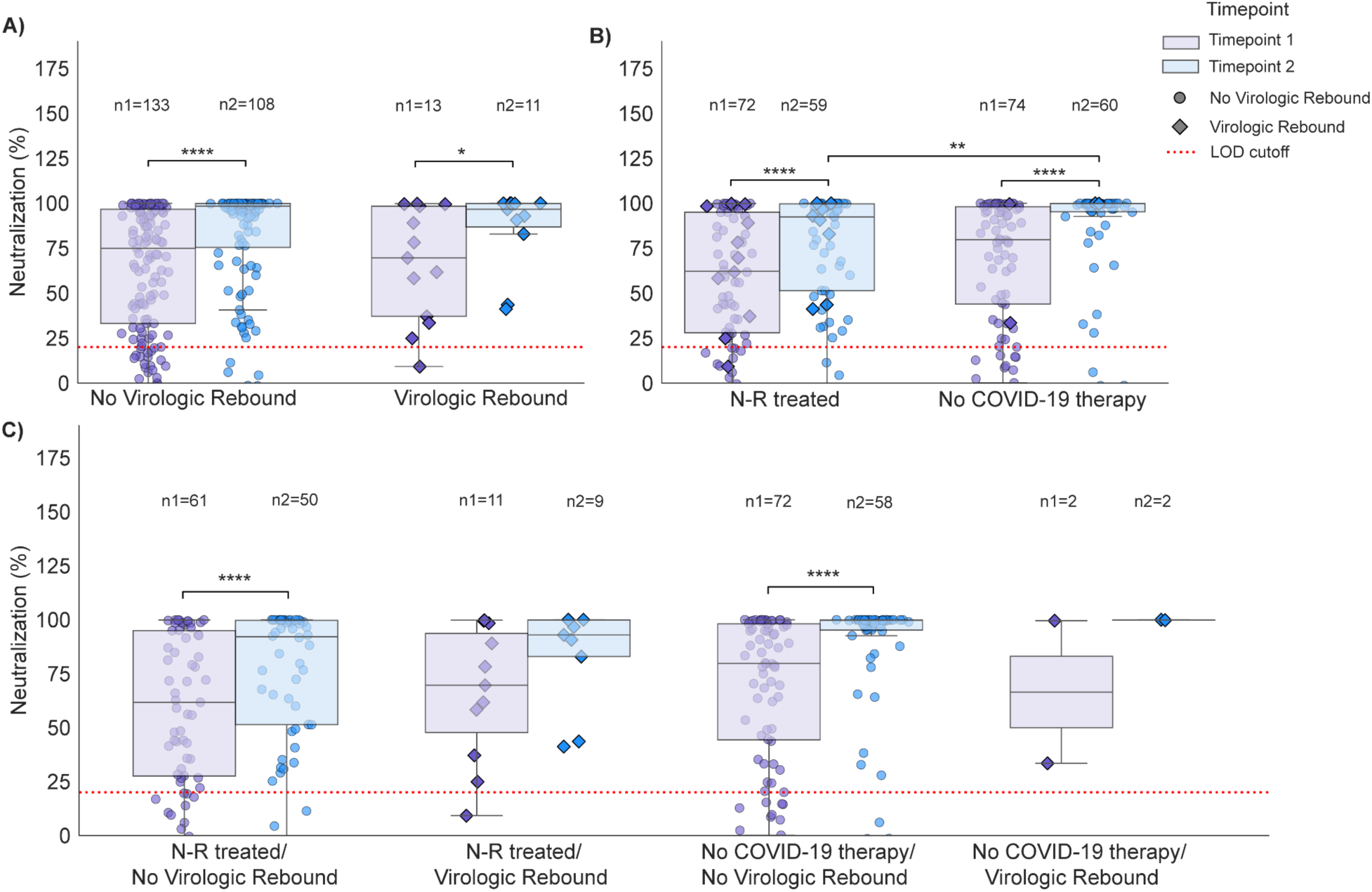
Neutralizing antibody levels against SARS-CoV-2 Omicron variant at Timepoints 1 and 2. Serum neutralization of SARS-CoV-2 B.1.1.529/Omicron in **A)** All participants stratified by virologic rebound status (no virologic rebound vs. virologic rebound). **B)** Participants stratified by treatment: N-R treated vs. no COVID-19 therapy. **C)** Participants stratified jointly by treatment and rebound status, yielding four groups: no virologic rebound / N-R treated, no virologic rebound / no COVID-19 therapy, virologic rebound / N-R treated, and viral rebound / no COVID-19 therapy. Timepoint 1 represents 0-15 days after detection (acute phase), Timepoint 2 represents 15 to 60 days after detection (post-acute phase). Sample sizes per timepoint (n1–n4) are indicated above each group. Multiple comparisons for different timepoints were performed using Wilcoxon matched-pairs signed rank test. Comparisons between independent groups were performed using the Mann-Whitney U test. Box plots show 25th, 50th, and 75th percentiles; whiskers, maximum and minimum. All data points are shown; y-axes show percentage. Diamond symbols represent virologic rebounders. Symbols above the brackets indicate the degree of significance. No asterisks is nonsignificant,**** p<0.0001, *** *P* < 0.001, ** *P* < 0.01 and * *P* < 0.05.

We examined neutralizing antibody responses against B.1.1.529 (Omicron), the infecting variant in all participants, in greater detail. Neutralizing antibody levels increased over time following acute infection, with significantly higher responses observed at Timepoint 2 compared with Timepoint 1 (Figure 2), consistent with the kinetics observed for binding antibody responses (Figure 1). This temporal increase was observed irrespective of COVID-19 treatment or virologic rebound status. When stratified by treatment status, individuals who did not receive COVID-19 therapy exhibited significantly higher neutralizing antibody levels at Timepoint 2 compared with individuals who received N-R treatment (p < 0.01) (Figure 2B).

After further stratification by virologic rebound status and COVID-19 treatment, the groups without virologic rebound demonstrated the expected longitudinal increase in neutralizing antibody responses, characterized by higher antibody levels at Timepoint 2 than at Timepoint 1. A similar trend was observed among N-R–treated individuals with virologic rebound (n at Timepoint 1 = 11; n at Timepoint 2 = 9) and untreated individuals with rebound (n at Timepoint 1 = 2; n at Timepoint 2 = 2); however, these differences did not reach statistical significance (p > 0.05), possibly due to the reduced power for these comparisons (Figure 2C). Consistent with the binding antibody findings, individuals with severe immunosuppression produced antibodies with negligible neutralizing activity (Figure S4).

To adjust for multiple variables that may affect neutralization responses, we performed multivariable analysis. Because of the strong ceiling effect observed in neutralization measurements, we performed logistic regression with an outcome variable separating near-complete neutralization (≥99%) from variation among sub-maximal responses (<99%). In logistic regression models, N-R treatment was associated with lower odds of achieving near-complete neutralization (OR = 0.41, p = 0.01), while the probability of near-complete neutralization increased significantly with time since infection (p < 0.001) (Table 3). Among samples with sub-maximal neutralization, neutralization also increased with time since infection (p <0.001), but was not significantly associated with N-R treatment or rebound status; however, it was associated with immunosuppression (Table 4).

**Table 3:** Multivariable logistic regression analysis of factors associated with high neutralization (≥99%).

| Variable | Odds Ratios (95%<br>CI) | *p-value* |
| --- | --- | --- |
| (Intercept) | 1.44 (0.92–2.29) | 0.113 |
| Immunosuppressed [Yes] | 0.72 (0.32–1.60) | 0.426 |
| Experienced virologic rebound [Yes] | 1.42 (0.40–5.11) | 0.585 |
| N-R [Yes] | 0.41 (0.20–0.81) | **0.011** |
| Days since infection (scaled) | 2.80 (1.67–4.90) | **<0.001** |

•OR = odds ratio
•CI = confidence interval
•Days since infection was standardized (mean-centered and scaled by SD)
•Reference groups: non-IS cohort, no viral rebound, no N-R

**Table 4:** Regression coefficients for the fitted linear mixed effects model for low neutralization.

| Predictors | Estimates (B) | SE | *p-value* |
| --- | --- | --- | --- |
| (Intercept) | 71.33 | 5.81 | *** |
| Immunosuppressed<br>[Yes] | -25.36 | 9.95 | * |
| Experienced virologic<br>rebound [Yes] | 10.98 | 16.32 | 0.504 |
| N-R [Yes] | 3.45 | 8.79 | 0.70 |
| Days since infection<br>(scaled) | 9.79 | 2.17 | *** |

Reference groups: Not immunosuppressed, did not experience SARS-CoV-2 viral rebound, did not receive N-R treatment
Symbols indicate the degree of significance. No asterisks is nonsignificant, and \*\*\* equates to p- value < 0.001

### Spike-specific T-cell responses are similar among subjects with and without virologic rebound

Given the central role of cellular immunity in viral clearance and control (33–35), we next measured Spike-specific T-cell responses by IFN-γ fluorospot assays. This analysis was performed in individuals for whom PBMC were available, a subset of participants (Total n=93, N-R treated n=43, No COVID-19 therapy n=50). IFN-γ producing T-cell responses were similar at Timepoint 1 and Timepoint 2 in N-R treated individuals compared to those individuals who did not receive any COVID-19 therapy (Figure 3A). Among N-R treated individuals, Spike-specific T-cell responses were slightly higher at Timepoint 1 compared to Timepoint 2 (Figure 3A). Individuals with virologic rebound showed higher IFN-γ producing units per million cells following Spike peptide stimulation compared to those without rebound, although numbers were small (n=6 in the rebound group) and the difference did not reach statistical significance (Figure 3B and C).

**Figure 3.**
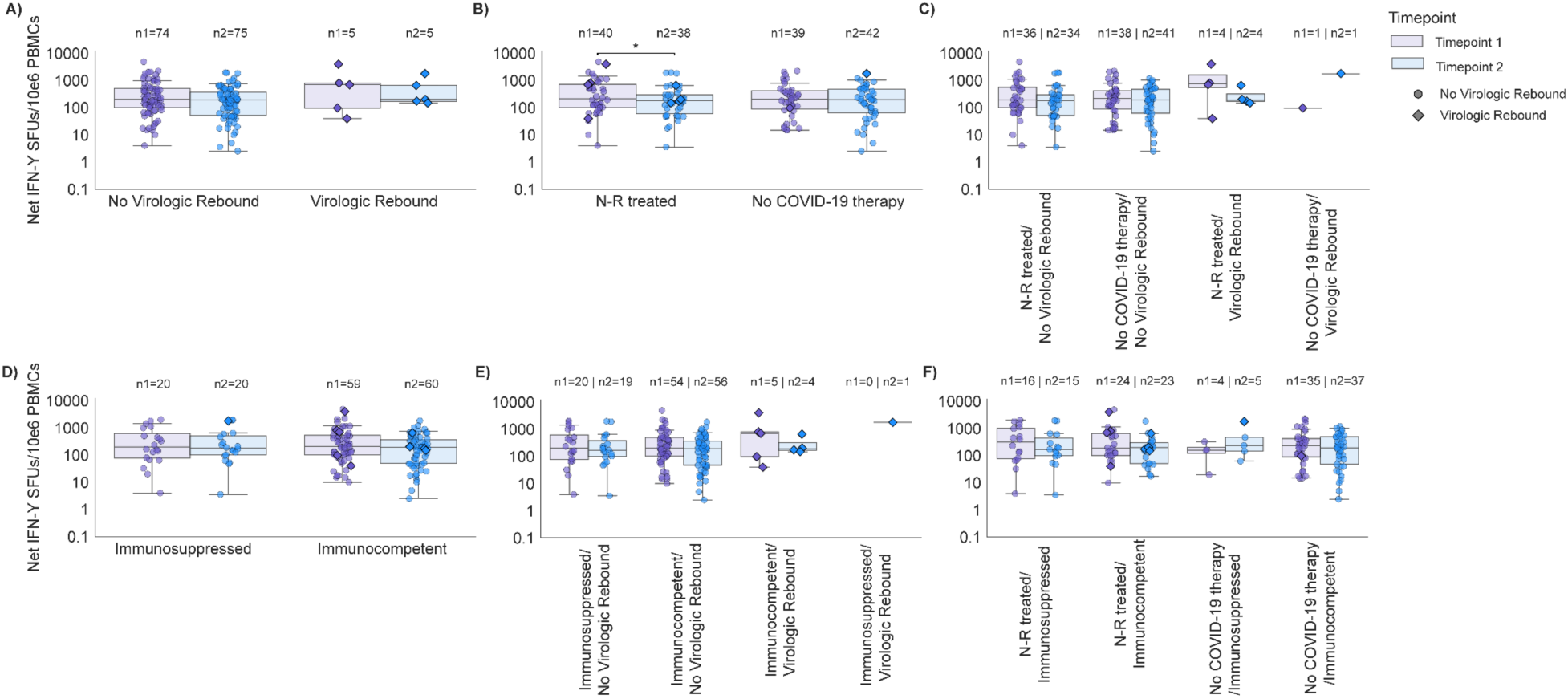
T-cell responses to SARS-CoV-2 antigen. Spike peptide pool–induced IFN-γ fluorospot responses are shown for **(A)** individuals with and without virologic rebound; **B)** N-R–treated individuals versus individuals without COVID-19 therapy; **(C)** stratified groups: N-R treated without virologic rebound, untreated without virologic rebound, N-R–treated with virologic rebound, and untreated without virologic rebound; **(D)** individuals with immunosuppressed and immunocompetent status; **E)** stratified groups: immunosuppressed without virologic rebound, immunocompetent with no virologic rebound, immunosuppressed with virologic rebound, and immunocompetent with virologic rebound; and **(F)** stratified groups: immunosuppressed N-R treated, immunocompetent N-R–treated, immunosuppressed untreated, and immunocompetent N-R–treated.. Responses were measured at Timepoint 1 (acute phase of infection) and Timepoint 2 (post-acute phase of infection). Pairwise comparisons between treatment and rebound groups were performed using Mann–Whitney tests, whereas Wilcoxon matched-pairs signed-rank tests were used for longitudinal analyses. Box plots show the 25th, 50th (median), and 75th percentiles; whiskers indicate the minimum and maximum values. All data points are shown. Y-axes are displayed on a logarithmic scale. Diamond symbols represent virologic rebounders. Symbols above the brackets indicate the degree of significance. No asterisks is nonsignificant,**** p<0.0001, *** *P* < 0.001, ** *P* < 0.01 and * *P* < 0.05.

When stratified by immune status, immunocompromised individuals showed numerically higher Spike-specific IFN-γ responses during acute phase of infection compared to immunocompetent individuals, although the difference was not statistically significant (Figure 3D). Additional stratification by immune status and virologic rebound similarly showed numerically higher Spike-specific IFN-γ responses among participants with virologic rebound without reaching statistical significance (Figure 3E). No significant differences in Spike-specific T-cell responses were observed when individuals were grouped by immune status and N-R treatment (Figure 3F).

We also analyzed T-cell responses using a linear mixed-effects model. No demographic or clinical variables were associated with Spike-specific cellular immune responses (Table 5). Point estimates suggest a possible increase in responses among participants with virologic rebound, but confidence intervals were wide and overlapped zero.

**Table 5:** Regression coefficients for the fitted linear mixed effects model for low T-cell immune responses.

| Predictors | Estimates<br>(B) | SE | *p-value* |
| --- | --- | --- | --- |
| (Intercept) | 2.23 | 0.09 | *** |
| Immunosuppressed<br>[Yes] | 0.06 | 0.14 | 0.688 |
| Experienced virologic<br>rebound [Yes] | 0.27 | 0.23 | 0.240 |
| N-R [Yes] | 0.10 | 0.12 | 0.410 |
| Days since infection | -0.003 | 0.003 | 0.289 |

Overall, anti-Spike T-cell responses in Timepoint 2 were comparable to those observed at Timepoint 1, indicating that robust Spike-specific T-cell immunity was already present during the acute phase of infection. This likely reflects high levels of pre-existing anti-Spike T-cell immunity within the cohort. Although limited longitudinal changes were observed, Spike-specific T-cell responses remained robust throughout the acute and post-acute phases of infection.

### Cytokine and chemokine levels are not associated with virologic rebound

Finally, we quantified the abundance of 25 cytokines and chemokines in subjects (n=36) with and without virologic rebound. We initially performed exploratory analyses across the complete cytokine and chemokine panel stratified by clinical variables and longitudinal timepoints. Detailed analyte-specific comparisons are provided in Supplementary data part 2, while Figure S5 and S6 highlight cytokines demonstrating the most notable temporal or treatment-associated patterns.

Overall, the analyses suggest that the evaluated clinical factors (virologic rebound status, N-R treatment or immunosuppressed status) were not associated with broad alterations in cytokine or chemokine concentrations, which remained relatively stable between Timepoint 1 and Timepoint 2, with only a few exceptions. Among the cytokines analyzed, IP-10, an interferon-inducible inflammatory chemokine, demonstrated the greatest magnitude of temporal and subgroup-associated variation across the cytokine panel (Figure S4 and Figure S5). IP-10 concentrations were generally higher at Timepoint 1 and decreased by Timepoint 2, suggesting resolution of interferon-associated inflammatory signaling during recovery from acute infection (36) (Figure S5). This decline was most evident among untreated, immunocompetent individuals and individuals who did not experience virologic rebound.

In contrast, only modest differences were observed for selected cytokines associated with inflammatory regulation and T-cell activation. MIP-1α and IL-2R concentrations were higher in N-R–treated individuals at Timepoint 1, whereas IL-1RA and IL-15 demonstrated modest differences between treatment groups at Timepoint 2 (Figure S6). However, these differences were not consistently associated with virologic rebound status.

To further visualize longitudinal cytokine patterns at the individual level, we generated paired-sample heatmaps using cytokine-wise Z-score normalization across all participants (Figure S4). Heatmap analysis demonstrated substantial inter-individual heterogeneity in cytokine and chemokine profiles; however, no distinct global cytokine signature clustered according to virologic rebound status, N-R treatment, or immune status. Most cytokines displayed relatively stable longitudinal patterns between acute and post-acute phases of infection, further supporting the absence of a systemic inflammatory profile associated with virologic rebound.

## DISCUSSION

In this longitudinal study, we assessed humoral, cellular, and innate immune responses that occur in response to acute SARS-CoV-2 infections with and without virologic rebound, stratifying analyses by N-R treatment and time. We found that immune responses among subjects with virologic rebound were similar to the immune responses observed without virologic rebound: antibody responses, particularly against Nucleocapsid antigens (where the confounding effect of vaccination is minimized), increased following acute disease, peaked in early convalescence, and waned thereafter. In comparing subjects with and without rebound, including in analyses stratified by N-R treatment status, responses among subjects with virologic rebound were generally indistinguishable from those without virologic rebound, although rebound without N-R treatment is exceedingly rare and responses in this subgroup were not well-characterized. Overall, we failed to identify immunologic differences between subjects with and without rebound.

In unadjusted and multivariate analyses, we observed slight reductions in levels of neutralizing antibodies associated with N-R treatment. We speculate that these reductions are due to reduced antigenic exposure following rapid viral suppression as a result of the drug rather than intrinsic impairment of B-cell function(24, 37, 38). Despite subtle quantitative differences, antibody kinetics appeared unaffected by N-R treatment or viral rebound status. As in previous studies(27, 39–41), severe impairment was observed only among individuals with profound immunosuppression, an effect that was independent of treatment status. Although the number of individuals in the group was small (n=5), the effects are similar to those observed in other cohorts(27, 42).

T-cell responses also did not differ between subjects with virologic rebound. The lack of differences extended to the analysis that was stratified by N-R treatment. In contrast to antibody responses, immunocompromising conditions did not markedly affect Spike-specific T-cell responses, consistent with observations elsewhere(27, 43). Functional T-cell integrity was further supported by stable levels of cytokines associated with adaptive immune polarization, which were predominantly low and not influenced by stratification. Cytokines involved in T-cell homeostasis and activation also demonstrated largely stable profiles with some transient elevations in specific subsets which normalized over time, suggesting temporary immune engagement rather than sustained dysregulation. The preserved humoral kinetics and intact spike-specific T-cell responses observed in our cohort and are aligned with previous results (24, 27, 37). Together, these findings suggest that N-R treatment does not substantially impair cell-mediated immune responses and that differences in T-cell responses are unlikely to explain the occurrence of virologic rebound.

Innate inflammatory signaling remained tightly regulated over time, with no evidence of progressive systemic cytokine amplification. Although chemokine such as IP-10 exhibited dynamic regulation, these changes were consistent with ongoing immune engagement and resolution rather than uncontrolled inflammation. Regulatory cytokines were intermittently elevated or showed inter-individual variability with no signature pattern associated with treatment status, immune status, or viral rebound suggesting preservation of immune regulatory balance rather than widespread inflammatory dysregulation (44).

Strengths of this study include comprehensive profiling of the immune responses in a well-characterized cohort of symptomatic outpatients with COVID-19 with systematic sampling over time. Limitations of the study include the non-random assignment of N-R, and the relative rarity of virological rebound, particularly in the untreated group. Furthermore, the study contained only mildly-ill outpatients. Previous studies have shown that mild COVID-19 cases have regulated cytokine/chemokine profiles and coordinated immune response, in contrast to inflammatory dysfunction in severe COVID-19 cases (36, 45, 46). It is possible that a study of more severely-ill subjects would have shown greater differences; however, virologic rebound tends to be clinically mild when it does occur(18, 19).

Together, these data indicate coordinated and controlled immune responses across innate, regulatory, and adaptive compartments, without evidence of treatment-induced immune impairment or a cytokine signature associated with treatment regimen/immune status/viral rebound. Our findings suggest that viral rebound following N-R treatment in mildly symptomatic COVID-19 is unlikely to result from impaired humoral, cellular, or innate immunity. Instead, rebound after N-R therapy may reflect non-immune mechanisms, including the preservation of the viral target cells with incomplete viral clearance among N-R treated subjects (21) and/or persistence of infectious SARS-CoV-2 despite inhibition of MPro(22), rather than an attenuated or exuberant human immune response.

## Methods

### Parent Study Design

Adult outpatients in the Mass General Brigham health care system in Boston, Massachusetts with acute COVID-19 were enrolled in the POSt-VaccInaTIon Viral CharactEristics Study (POSITIVES) prospective cohort study beginning in 2022. All participants provided informed consent^14^. The medical records of the consenting participants were reviewed by study board-certified physicians for extraction of demographic data, medical history, COVID-19 vaccination status, and treatment history. Participants self-collected anterior nasal swabs every two to three days for a total of five to six samples over 2 weeks. Participants in POSITIVES were followed until they had two consecutive negative PCR tests. Blood collection was offered in a sub-study. Those who consented to this sub-study had blood collected at 4 timepoints after enrollment ^14,15,20,25–29^.

### Eligibility Criteria

For this analysis, we analyzed data from ambulatory patients in the POSITIVES study with acute COVID-19 enrolled on or after March 1, 2022. Participants were excluded based on the following criteria: 1) had no viral load and culture results at least 12 days after the initial diagnostic test, 2) a first nasal swab before March 2022 (when N-R became available), 3) inpatient, 4) N-R treated participants with a first study swab more than one day after N-R regimen completion, 6) participants who received co-therapy (remdesivir or molnupiravir) within the past two weeks or monoclonal antibodies past 90 days or non-standard N-R regimen and 7) no blood collected as part of the study.

### Study approval

Study procedures were reviewed and approved by the Human Participants IRB at Mass General Brigham under protocol no. 2021P000812.

### Sample collection and processing

Phlebotomy was performed at 4 different time points after SARS-CoV-2 was detected: Timepoint 1 was collected as early as possible after enrollment and before day 15 since symptoms onset or first positive PCR or antigen test (acute phase). Timepoint 2 was collected 15 to 60 days after COVID-19 detection (convalescent phase). Timepoint 3 was collected after 6 months, and Timepoint 4 was collected 12 months after the index episode of COVID-19. Blood was collected in sodium heparin BD cell preparation tubes and silica red top tubes (serum). Serum and PBMCs (BD cell preparation tubes) were isolated within 4 hours of collection. PBMC isolation was performed according to the manufacturer’s protocol for cell preparation tubes. Sera was stored at −80 degrees Celsius and PBMCs were stored in liquid nitrogen.

### Categorization for Immunosuppressed state

The categorization for immunosuppressed state has been explained in detail in our previous study^27^. Immunosuppressed participants were further categorized as severe (consisting of hematological malignancy/transplant participants and participants with autoimmune condition receiving B-cell targeting agents or B-cell deficiency) and non-severe^27^.

### Luminex assay

To measure concentrations of anti-Spike and anti-Nucleocapsid antibodies, a luminex assay kit from Invitrogen, ProcartaPlex™ Human Coronavirus Ig Total Panel, 11plex (Catalog number EPX110-16000-901) was used. Serum samples were diluted 1:1000-1:100,000 and processed as recommended by the kit and run through Luminex xMAP Intelliflex®. The concentrations of the antibodies were calculated using the Procartaplex analysis application. To analyze the variant specific neutralization potential of serum, the SARS-CoV-2 Variants Neutralizing Antibody 6-Plex ProcartaPlex™ Panel ( Invitrogen, Catalog number EPX060-16018-901) was used. Briefly, samples were diluted 1:100 and the assay was set up according to the manufacturer’s recommendations and run on the Luminex xMAP Intelliflex® system. Percent neutralization values were calculated from mean fluorescence values according to kit instructions.

To quantify cytokines and chemokines in the serum samples, the Cytokine 25-plex human panel luminex assay kit from Invitrogen (Catalog number LHC0009M) was used. The protocol provided by the manufacturer was once again followed; as per recommendation, the final dilution of serum was kept at 1:2 and run on the Luminex xMAP Intelliflex® system along with the standards given in the kit. The standard curve was made from the using a 5PL nonlinear regression model from the measurements obtained for the serially diluted standards and the concentrations of the serum samples were interpolated using GraphPad Prism (Version 10.6.1). Concentrations which were below the lower limit of quantification (LLOQ) were reported as LLOQ/2 as recommended by the manufacturer and multiplied by dilution factor. The concentrations calculated were transformed to a log10 scale before further analyses.

### Ex-vivo Fluorospot assay

An ex-vivo IFN-gamma fluorospot assay was performed to quantify the Spike-specific T-cell responses. Fluorospot plates from Mabtech pre-coated with anti-human IFN-ɣ monoclonal antibodies (clone 1-D1K) were used according to the manufacturer’s instructions (Mabtech) and as previously described^27^. Briefly, 100,000 to 200,000 peripheral blood mononuclear cells (PBMCs) were seeded per well to generate a monolayer and stimulated with 1 μg/ml SARS-CoV-2 peptide pools (JPT Peptide Technologies PM-WCPV-S-1, PepMix™ SARS-CoV-2 (Spike Glycoprotein)) for 16 to 18 hours at 37°C. Anti-CD3 mAb (CD3-2) and Anti-CD28 mAb (CD28-A) were used as positive controls and dimethyl sulfoxide (DMSO) as the negative control. Following incubation, spots were enumerated using an automated Fluorospot reader (MABtech IRIS 2). Antigen-specific responses were calculated by subtracting mean spots of the DMSO control wells from the peptide-stimulated wells and reported as fluorospot-forming units (FFU) per 10^6^ PBMCs. Responses were considered positive if the results were >5 FFU/10^6 PBMCs following background subtraction ^27^. If DMSO control wells had >30 FFU/10^6 PBMCs or if positive control wells (anti-CD3/anti-CD28 stimulation) were negative, the results were excluded from further analysis ^27^.

### Statistics

Categorical variables of the cohort were summarized using the total number and percentages, and a chi-square test or Fisher’s exact test was used to check for significant differences. Continuous variables were summarized with median and interquartile ranges, presented as box plots and compared with non-parametric methods to determine significance: either Wilcoxon matched-pairs signed rank test or Multiple Mann-Whitney tests or Kruskal–Wallis test. Statistical analyses and figure production were done using R (R Foundation for Statistical Computing, version 2024.04.2+764), GraphPad Prism, version 10 and Python. The lme4 package^30^ was used to create mixed effects linear regression models (GLMM) to estimate the effect of time, N-R receipt, sex, age and the vaccination status on SARS-CoV-2 Nucleocapsid titer and Spike-specific FFUs/million PBMCs. A GLMM was selected to allow us to incorporate random effects, therefore accounting for repeated measurements of participants. An interaction term was included between time and participant type for the antibody model. Neutralization antibody responses exhibited a strong ceiling effect, with nearly half of measurements at ≥99% neutralization, precluding analysis using standard Gaussian or beta regression models. We therefore analyzed neutralization using a two-part modelling strategy. First, the probability of near-complete neutralization (≥99%) was modeled as a binary outcome using logistic regression, adjusted for the same covariates listed above. Mixed-effects logistic models with a participant-level random intercepts were explored but not identifiable because of sparse repeated measurements (one to two timepoints per participants) and limited within-participant variation, so fixed-effects logistic regression was ultimately used. Among samples with sub-maximal neutralization (<99%), neutralization was analyzed as a continuous outcome using linear mixed-effects models with a participant-level intercept to account for repeated measurements. ChatGPT (OpenAI) was used to assist with code for plotting data. All analyses and outputs were independently verified by the authors.

## Supporting information

Supplementary data part 1

Supplementary data part 2

## Acknowledgments

The study team would like to thank all the participants for their participation in this study and all the healthcare workers who take care of them and kindly refer them to the POSITIVES study. This work was supported by the National Institutes of Health (NIH) (R01 AI 138801), the Massachusetts Consortium on Pathogen Readiness, and the Massachusetts General Hospital. Chloe M.Hasund was supported by T32 institutional training award from NIAID/NIH (T32AI049928 to Dyann Wirth).

## Author contributions

Conceptualization and study design: J.E.L, M.J.S., A.K.B. and J.Z.L. Data collection: D.G., J.B.C., M.Y.L., E.A, M.B., A.A., Z.R., E.P., C.M., Z.S.S., K.S., J.B., O.T.G., B.L., G.E.E., M.C.C.,Y.L.,N.P., J.E.S.. Analysis and data interpretation: D.G., C.M.H., M.Y.L., B.L.K., J.B., G.E.E., M.C., Y.L., J.E.S., J.E.L., M.J.S. Statistical analysis: D.G., C.M.H., B.L.K. Funding acquisition: J.E.L., M.J.S., A.K.B., J.Z.L. Writing the initial draft: D.G., J.E.L., C.M.H. Critical review of the manuscript and final draft: All authors.

## Notes

Conflict of Interest: JZL is supported by a grant from the Investigator-Initiated Studies Program of Merck Sharp & Dohme LLC. The opinions expressed in this paper are those of the authors and do not necessarily reflect those of Merck Sharp & Dohme LLC. YL is a topic editor for DynaMed All other authors declare no competing interests. JEL has received research grants from Moderna and Pfizer unrelated to this work. All other authors report no potential conflicts.

### Competing Interest Statement

JZL is supported by a grant from the Investigator-Initiated Studies Program of Merck Sharp & Dohme LLC. The opinions expressed in this paper are those of the authors and do not necessarily reflect those of Merck Sharp & Dohme LLC. YL is a topic editor for DynaMed All other authors declare no competing interests. JEL has received research grants from Moderna and Pfizer unrelated to this work. All other authors report no potential conflicts.

