## Supplementary figures for "Post-infection immune response in adults with COVID-19 with and without nirmatrelvir-ritonavir treatment and virologic rebound"

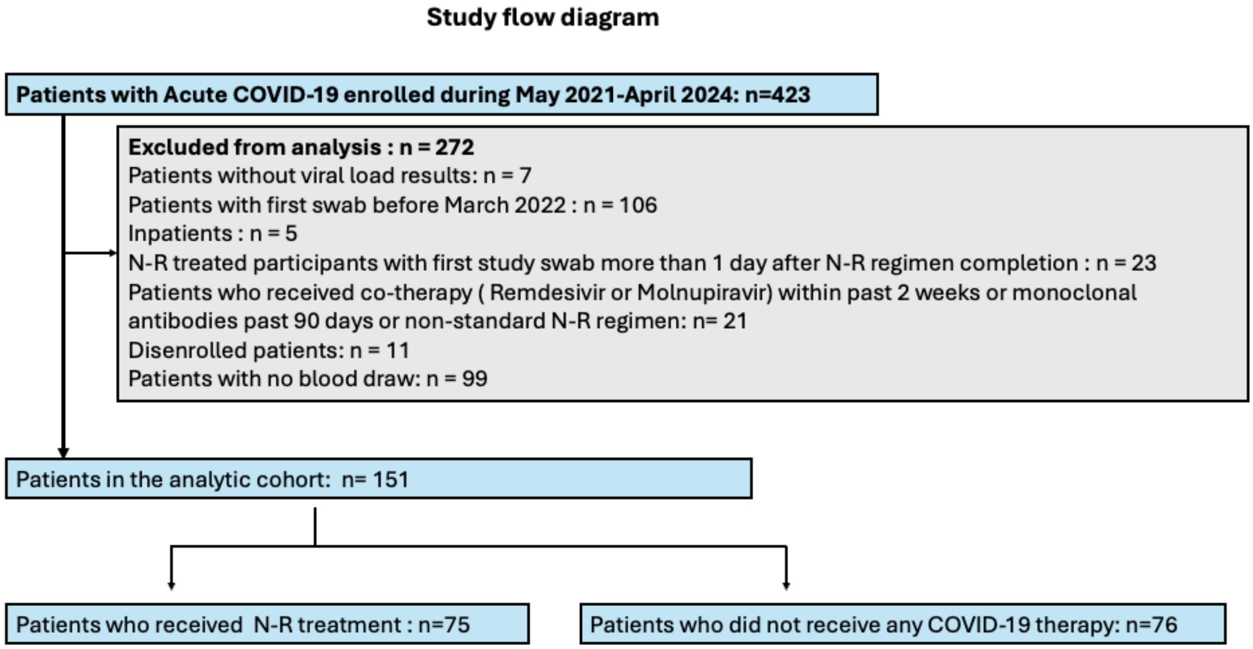


**Figure S1:** Flow diagram describing eligibility and enrollment criteria for this study.

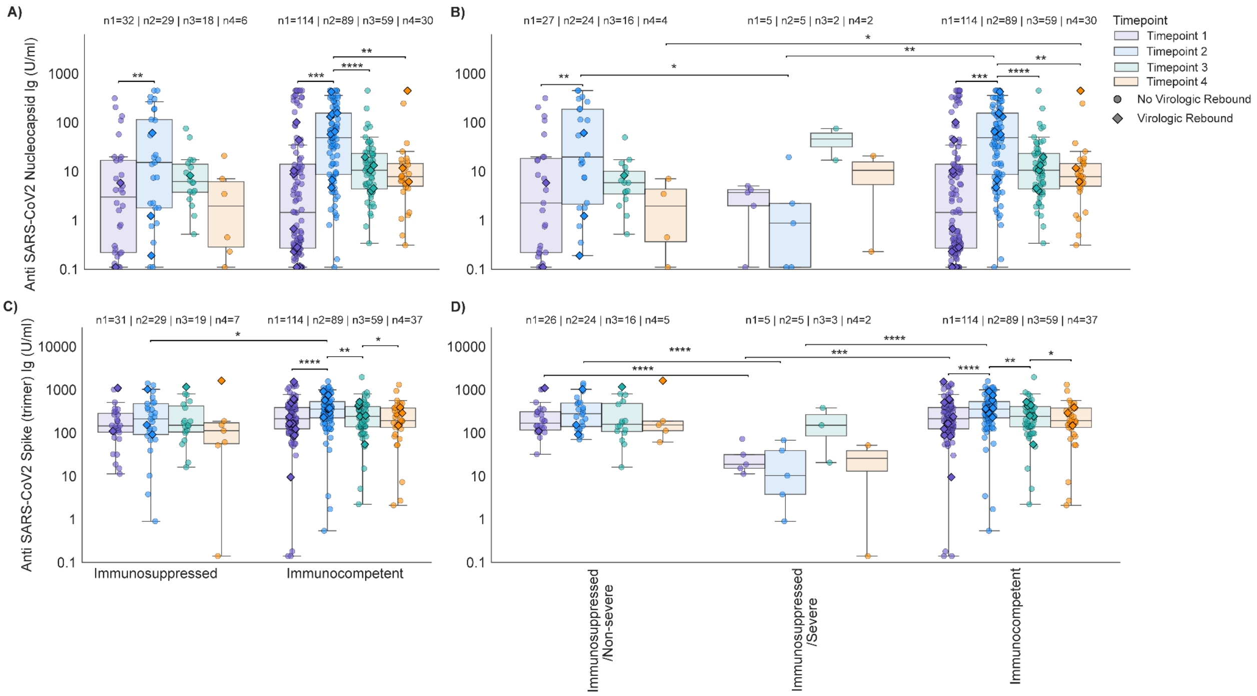


**Figure S2: Concentrations of binding antibodies over time in immunocompetent and immunosuppressed subjects.** Anti–SARS-CoV-2 Nucleocapsid **(A–B)** and Anti–SARS-CoV-2 Spike Trimer **(C–D)** antibody concentrations (U/mL) measured at four timepoints after COVID-19 detection. **(A, C)** All COVID-19 participantss stratified by Immune status (Immunosuppressed vs Immunocompetent). **(B, D)** Immunosuppressed participants were further subdivided into Severe and Non-severe based on immunosuppression severity. Timepoint 1 represents the acute phase (0–15 days after detection), Timepoint 2 the post-acute phase (15–60 days), Timepoint 3 approximately 6 months, and Timepoint 4 approximately 1 year after COVID-19 detection. Sample sizes per timepoint (n1–n4) are indicated above each group. Multiple comparisons across timepoints within a group were performed using the Wilcoxon matched-pairs signed rank test with Benjamini-Hochberg false discovery rate (FDR) correction. Comparisons between independent groups were performed using the Mann-Whitney U test. Box plots show the 25th, 50th, and 75th percentiles; whiskers indicate maximum and minimum values. All data points are shown; y-axes are on a logarithmic scale. Diamond symbols represent virologic rebounders. Symbols above the brackets indicate the degree of significance. No asterisks is nonsignificant,**** p<0.0001, *** $P < 0.001$, ** $P < 0.01$ and * $P < 0.05$.

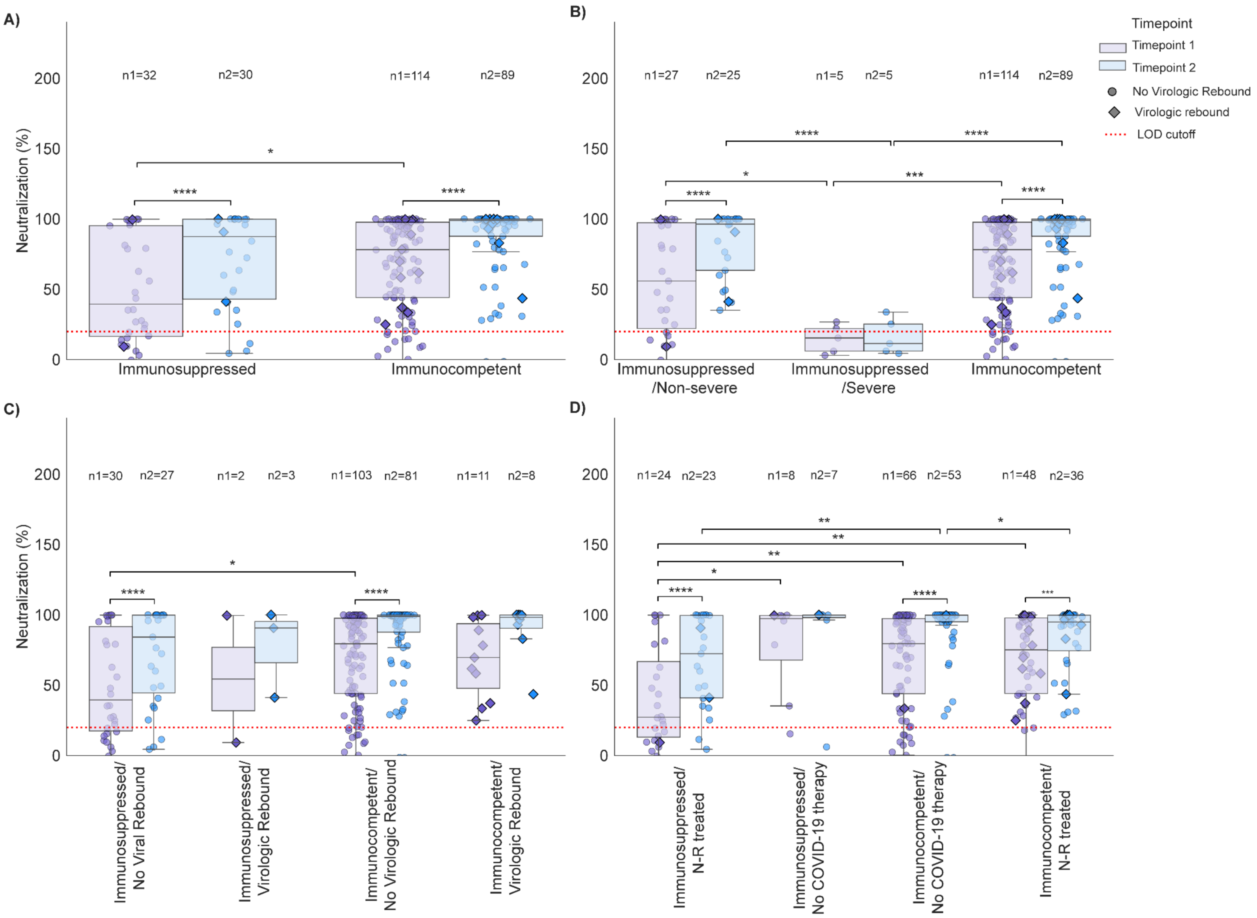


**Figure S3: Neutralizing antibody levels against SARS-CoV-2 B.1.1.529/Omicron over time. in Immunocompetent and Immunosuppressed participants.** Serum neutralization of in **A**) Immunocompetent and immunosuppressed individuals **B**) Immunocompetent and immunosuppressed individuals (with severe and non-severe immunosuppression) **C**) stratified groups: immunocompetent with virologic rebound, and immunocompetent with no virologic rebound, immunosuppressed with virologic rebound, immunosuppressed with no virologic rebound,; and **(D)** stratified groups: immunosuppressed N-R treated, immunocompetent untreated, immunocompetent untreated, and immunocompetent N-R–treated in 2 different timepoints after the detection of COVID-19. Timepoint 1 represents 0-15 days after detection (acute phase) and Timepoint 2 represents 15 to 60 days after detection (post-acute phase). Sample sizes per timepoint (n1–n2) are indicated above each group. Multiple comparisons for different timepoints were performed using Wilcoxon matched-pairs signed rank test. Comparisons between independent groups were performed using the Mann-Whitney U test. Box plots show 25th, 50th, and 75th percentiles; whiskers, maximum and minimum. All data points are shown; y-axes show percentage. The diamond dots represent virologic rebounders. Symbols above the brackets indicate the degree of significance. No asterisks is nonsignificant,**** p<0.0001, *** $P < 0.001$, ** $P < 0.01$ and * $P < 0.05$.

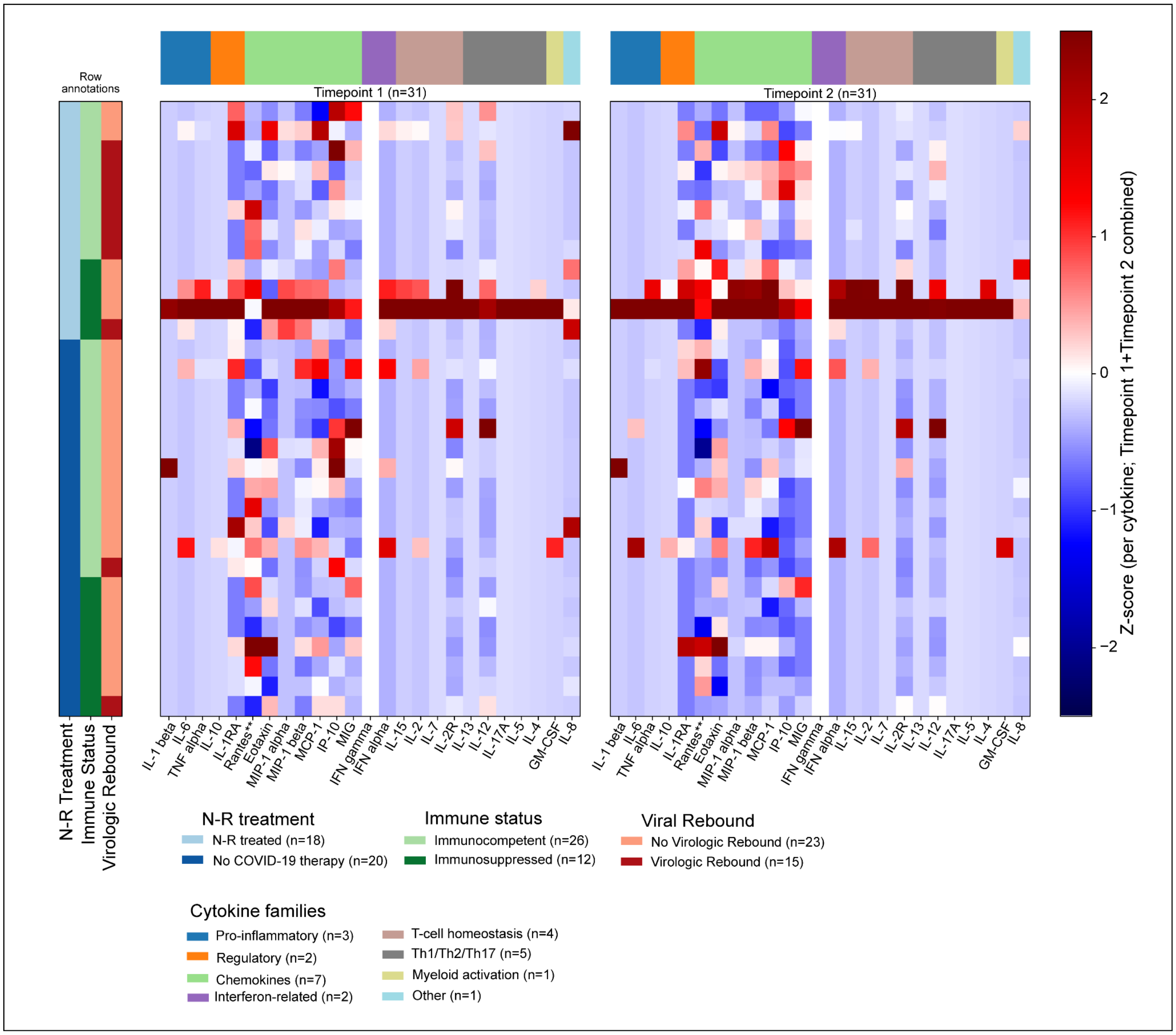


**Figure S4: Heatmap of cytokine and chemokine profiles across study groups.**
Heatmap of Z-score–normalized cytokine and chemokine levels measured by 25-plex Luminex assay across study participants. Values are Z-score–normalized within each cytokine across all participants, emphasizing relative differences between individuals for each analyte. The color intensity scale represents the z-score ranging from the highest (Red) to the lowest (dark purple). Samples were collected at Timepoint 1 (0-15 days after detection/ acute phase) and Timepoint 2 (15 to 60 days after detection/ post-acute phase) and heatmaps for both timepoints are displayed side-by-side using an identical color scale to enable direct comparison across timepoints. Rows represent individuals and presented in a predefined order and are not hierarchically clustered by clinical strata: Treatment (N-R treated and No COVID-19 therapy), then immune status (immunocompetent and immunosuppressed) and then virologic rebound status (No virologic rebound and virologic rebound) which is indicated by colored row-annotation bars. Columns represent cytokines/chemokines  measured and are grouped by functional family, with a column annotation bar indicating families (e.g., pro-inflammatory, chemokines, interferon-related, T-cell homeostasis, and other specified groups). Group sizes (n) for row annotation categories, column annotation categories and total samples per timepoint are shown on the figure.

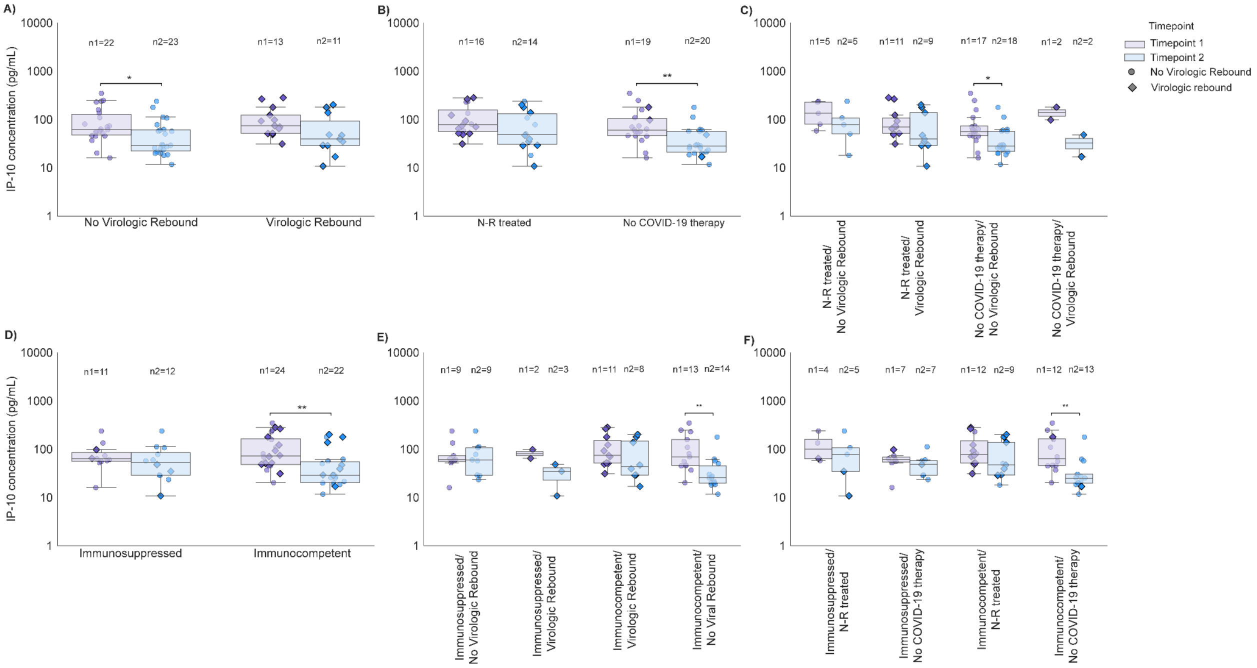


**Figure S5. Chemokine IP-10 dynamics over the time of COVID-19 infection.** Concentration (pg/ml) of IP-10 is shown for **(A)** individuals with and without virologic rebound; **B)** N-R–treated individuals versus individuals without COVID-19 therapy; **(C)** stratified groups: N-R treated without virologic rebound, untreated without virologic rebound, N-R–treated with virologic rebound, and untreated without virologic rebound; **(D)** individuals with immunosuppressed and immunocompetent status; **E)** stratified groups: immunosuppressed without no virologic rebound, immunocompetent with no virologic rebound, immunocompromised with rebound, and immunocompetent with virologic rebound; and **(F)** stratified groups: immunosuppressed N-R treated, immunocompetent N-R–treated, immunocompetent untreated, and immunocompetent N-R–treated.Responses were measured at Timepoint 1 (acute phase of infection) and Timepoint 2 (post-acute phase of infection). Sample sizes per timepoint (n1–n2) are indicated above each group. Pairwise comparisons between treatment and rebound groups were performed using Mann–Whitney tests, whereas Wilcoxon matched-pairs signed-rank tests were used for longitudinal analyses. Box plots show the 25th, 50th (median), and 75th percentiles; whiskers indicate the minimum and maximum values. All data points are shown. Y-axes are displayed on a logarithmic scale.The diamond dots represent virologic rebounders. Symbols above the brackets indicate the degree of significance. No asterisks is nonsignificant,**** p<0.0001, *** $P < 0.001$, ** $P < 0.01$ and * $P < 0.05$.

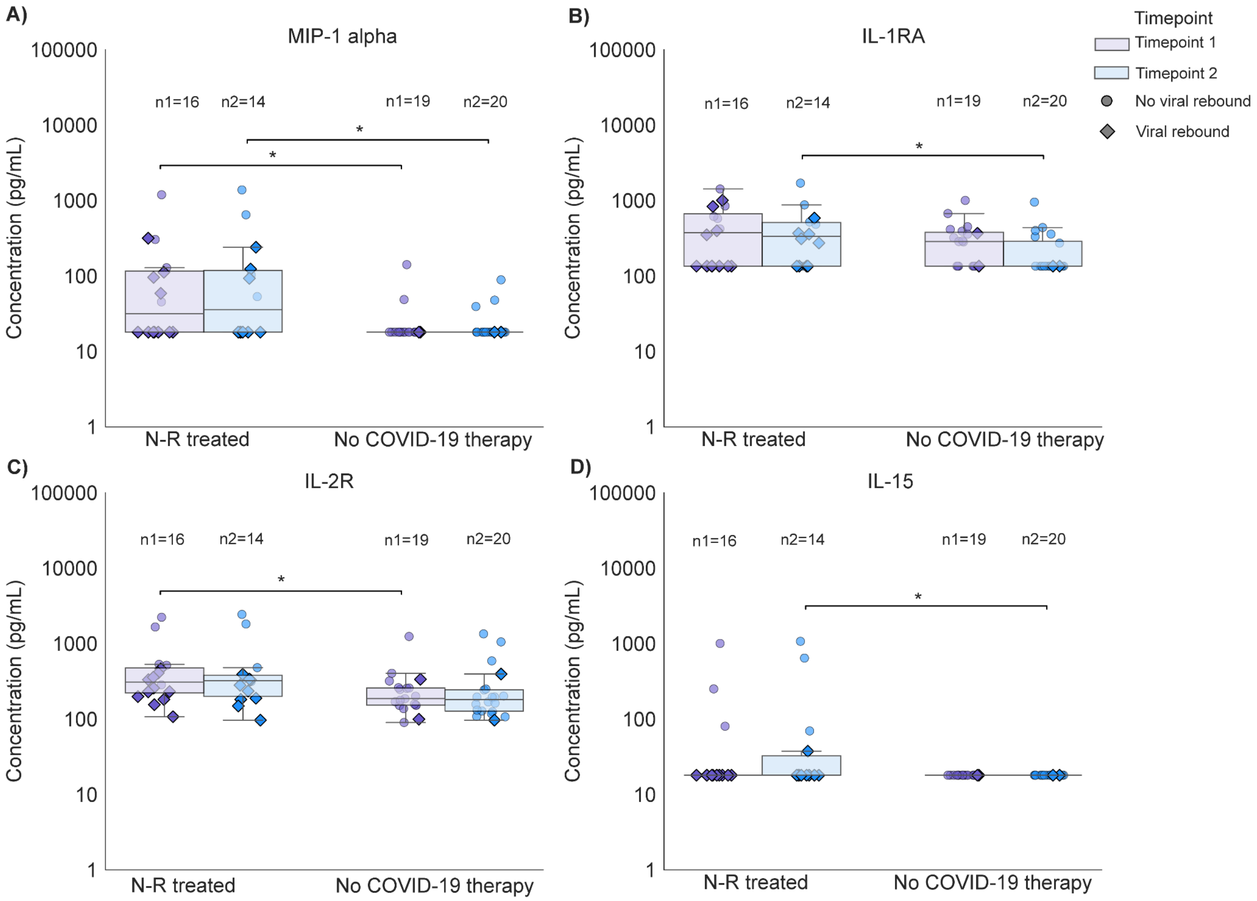


**Figure S6. Concentrations of chemokines/cytokines over time in N-R treated participants and participants who did not receive any COVID-19 therapy.** Concentrations of **A)** MIP-1 alpha, **B)** IL-1RA, **C)** IL-2R and **D)** IL-15 in N-R treated participants and participants who did not receive any COVID-19 therapy. Timepoint 1 represents the acute phase (0–15 days after detection) and Timepoint 2 represents the post-acute phase (15–60 days) after COVID-19 detection. Sample sizes per timepoint (n1–n2) are indicated above each group. Multiple comparisons across timepoints within a group were performed using the Wilcoxon matched-pairs signed rank test. Comparisons between independent groups were performed using the Mann-Whitney U test. Box plots show the 25th, 50th, and 75th percentiles; whiskers indicate maximum and minimum values. The diamond dots represent virologic rebounders. Symbols above the brackets indicate the degree of significance. No asterisks is nonsignificant,**** p<0.0001, *** $P < 0.001$, ** $P < 0.01$ and * $P < 0.05$.
