## Supplementary data part 1 for "Post-infection immune response in adults with COVID-19 with and without nirmatrelvir-ritonavir treatment and virologic rebound"

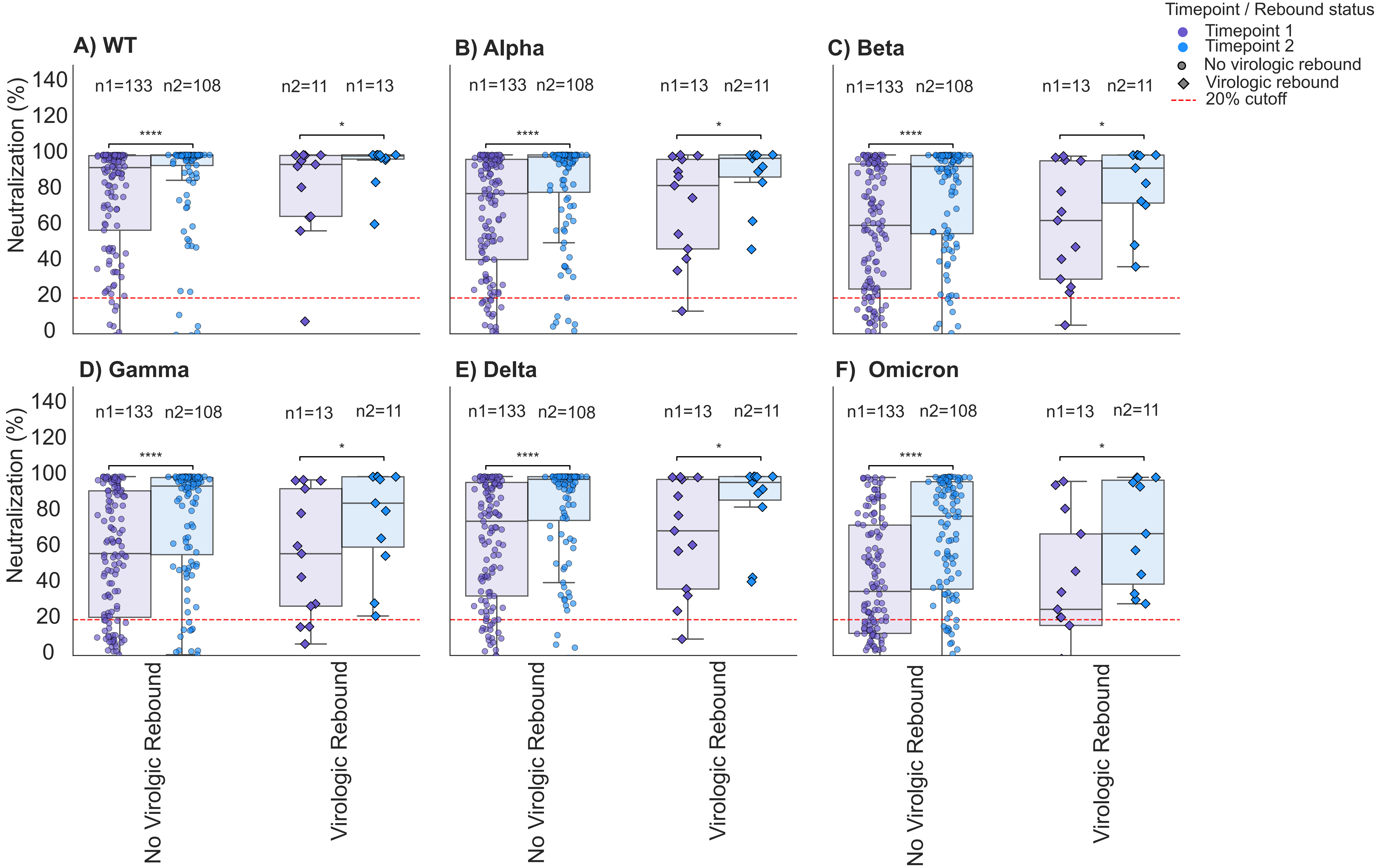


**Figure: Neutralizing antibody levels over time in participants who experienced viral rebound and who did not.** Serum neutralization of different variants of SARS-CoV-2 A) SARS-CoV-2 WT, B) SARS-CoV-2 B.1.1.7/ Alpha, C) SARS-CoV-2 B.1.351/Beta, D) SARS-CoV-2 P.1/Gamma, E) SARS-CoV-2 B.1.617.2/Delta and F) SARS-CoV-2 B.1.1.529/Omicron in Paxlovid treated and untreated participants in 2 different timepoints after the detection of COVID-19. Timepoint 1 represents 0-15 days after detection ( acute phase), Timepoint 2 represents 15 to 60 days after detection (post-acute phase). Sample sizes per timepoint (n1, n2) are indicated above each group. Multiple comparisons for different timepoints were performed using Wilcoxon matched-pairs signed rank test. Comparisons between independent groups were performed using the Mann-Whitney U test. Box plots show 25th, 50th, and 75th percentiles; whiskers, maximum and minimum. All data points are shown; y-axes show percentage. The diamond dots represent virologic rebounders. Symbols above the brackets indicate the degree of significance. No asterisks is nonsignificant, **** p<0.0001, *** P < 0.001, ** P < 0.01 and * P < 0.05.

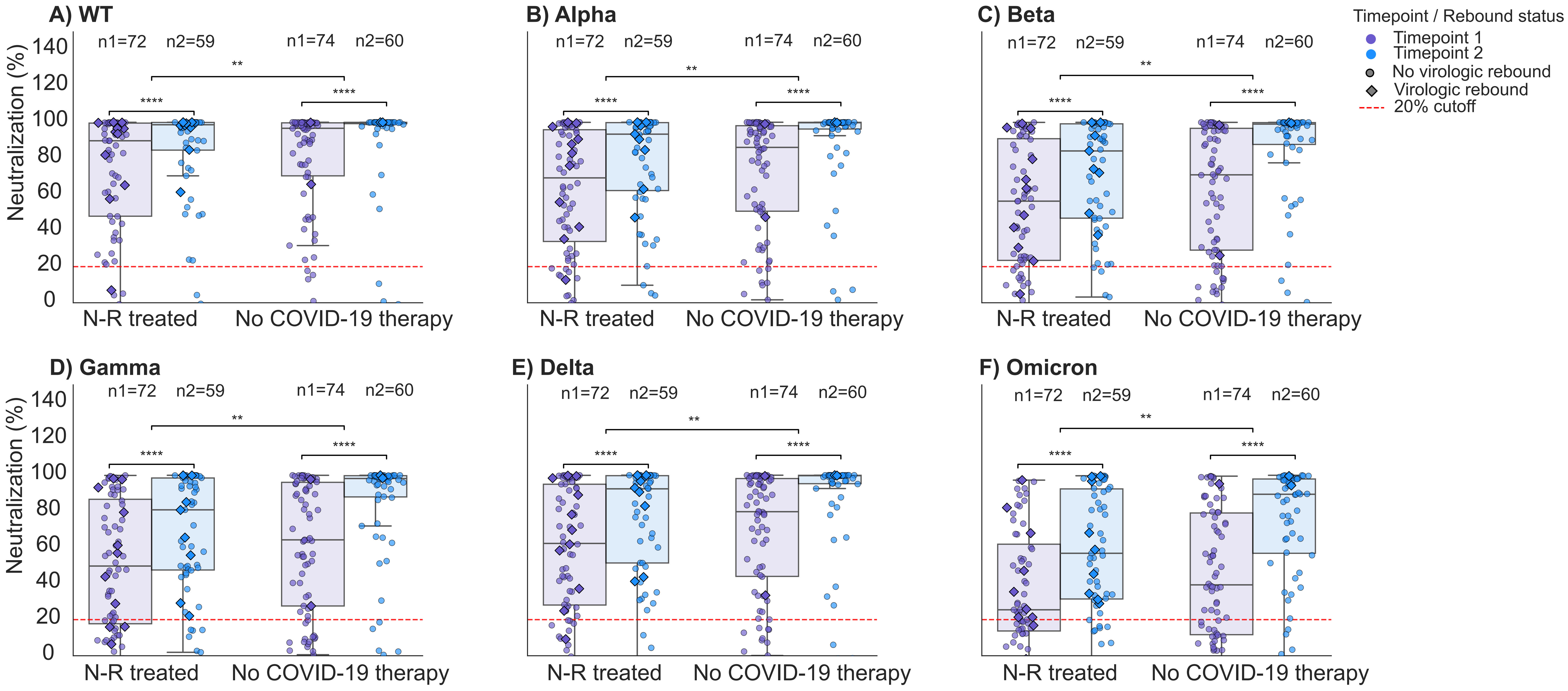


**Figure: Neutralizing antibody levels over time in participants who received N-R treatment and who did not receive any COVID-19 therapy.** Serum neutralization ofdifferent variants of SARS-CoV-2 A) SARS-CoV-2 WT, B) SARS-CoV-2 B.1.1.7/ Alpha, C) SARS-CoV-2 B.1.351/Beta, D) SARS-CoV-2 P.1/Gamma, E) SARS-CoV-2 B.1.617.2/Delta and F) SARS-CoV-2 B.1.1.529/Omicron in N-R treated and untreated participants in 2 different timepoints after the detection of COVID-19. Timepoint 1 represents 0-15 days after detection ( acute phase), Timepoint 2 represents 15 to 60 days after detection (post-acute phase). Sample sizes per timepoint (n1, n2 are indicated above each group. Multiple comparisons for different timepoints were performed using Wilcoxon matched-pairs signed rank test. Comparisons between independent groups were performed using the Mann-Whitney U test. Box plots show 25th, 50th, and 75th percentiles; whiskers, maximum and minimum. All data points are shown; y-axes show percentage. The diamond dots represent virologic rebounders. Symbols above the brackets indicate the degree of significance. No asterisks is nonsignificant, **** $p<0.0001$, *** $P < 0.001$, ** $P < 0.01$ and * $P < 0.05$.

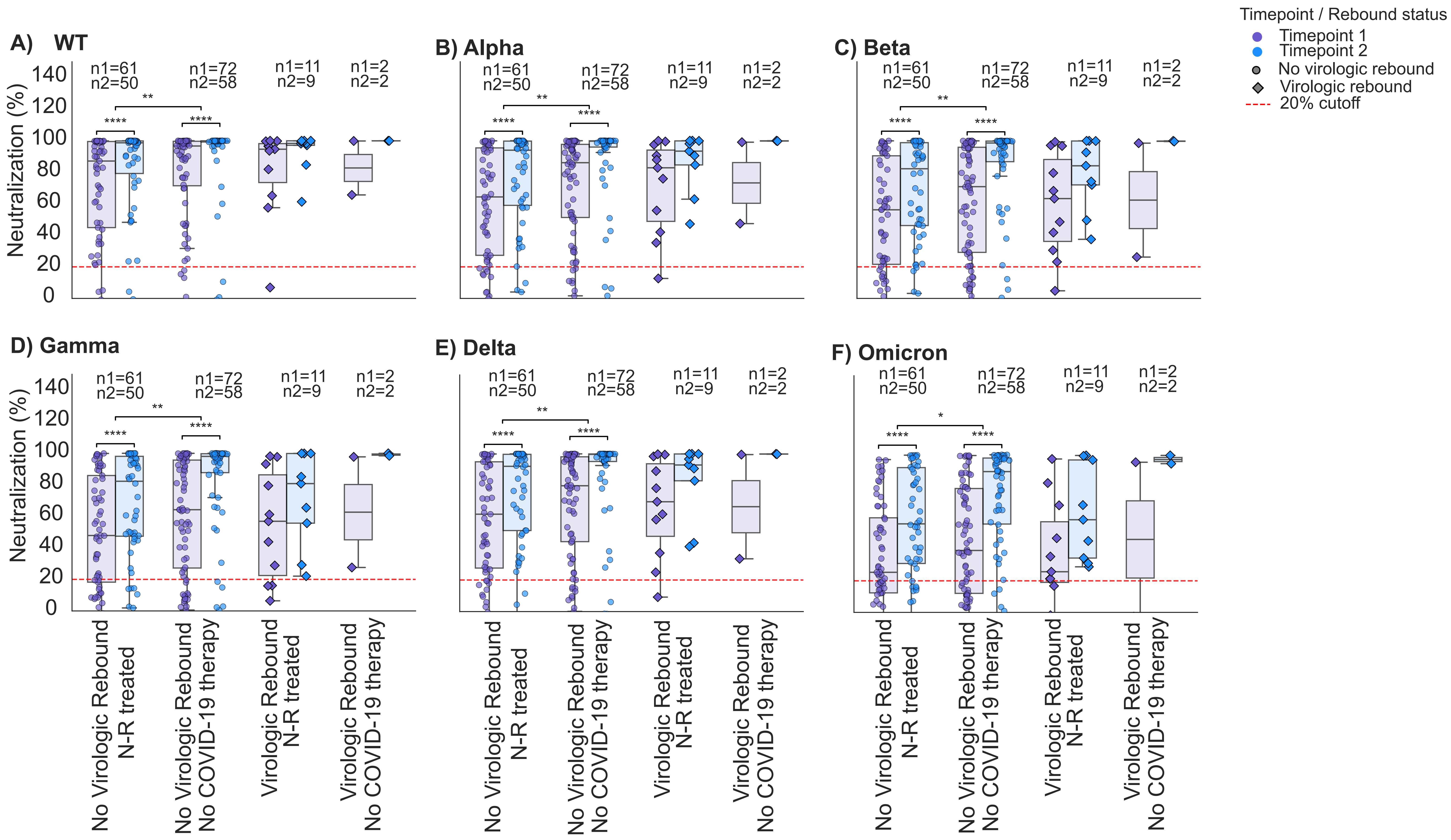


**Figure: Neutralizing antibody levels over time in participants in stratified groups: No viral rebound participants treated with N-R, No Viral rebound participants untreated with any COVID-19 therapy, Viral rebound experienced N-R treated participants and Viral rebound experienced participants who did not receive any COVID-19 therapy.** Serum neutralization ofdifferent variants of SARS-CoV-2 A) SARS-CoV-2 WT, B) SARS-CoV-2 B.1.1.7/ Alpha, C) SARS-CoV-2 B.1.351/Beta, D) SARS-CoV-2 P.1/Gamma, E) SARS-CoV-2 B.1.617.2/Delta and F) SARS-CoV-2 B.1.1.529/Omicron in the participants in 2 different timepoints after the detection of COVID-19. Timepoint 1 represents 0-15 days after detection ( acute phase), Timepoint 2 represents 15 to 60 days after detection (post-acute phase). Sample sizes per timepoint (n1, n2) are indicated above each group. Multiple comparisons for different timepoints were performed using Wilcoxon matched-pairs signed rank test. Comparisons between independent groups were performed using the Mann-Whitney U test. Box plots show 25th, 50th, and 75th percentiles; whiskers, maximum and minimum. All data points are shown; y-axes show percentage. The diamond dots represent virologic rebounders. Symbols above the brackets indicate the degree of significance. No asterisks is nonsignificant, **** p<0.0001, *** P < 0.001, ** P < 0.01 and * P < 0.05.

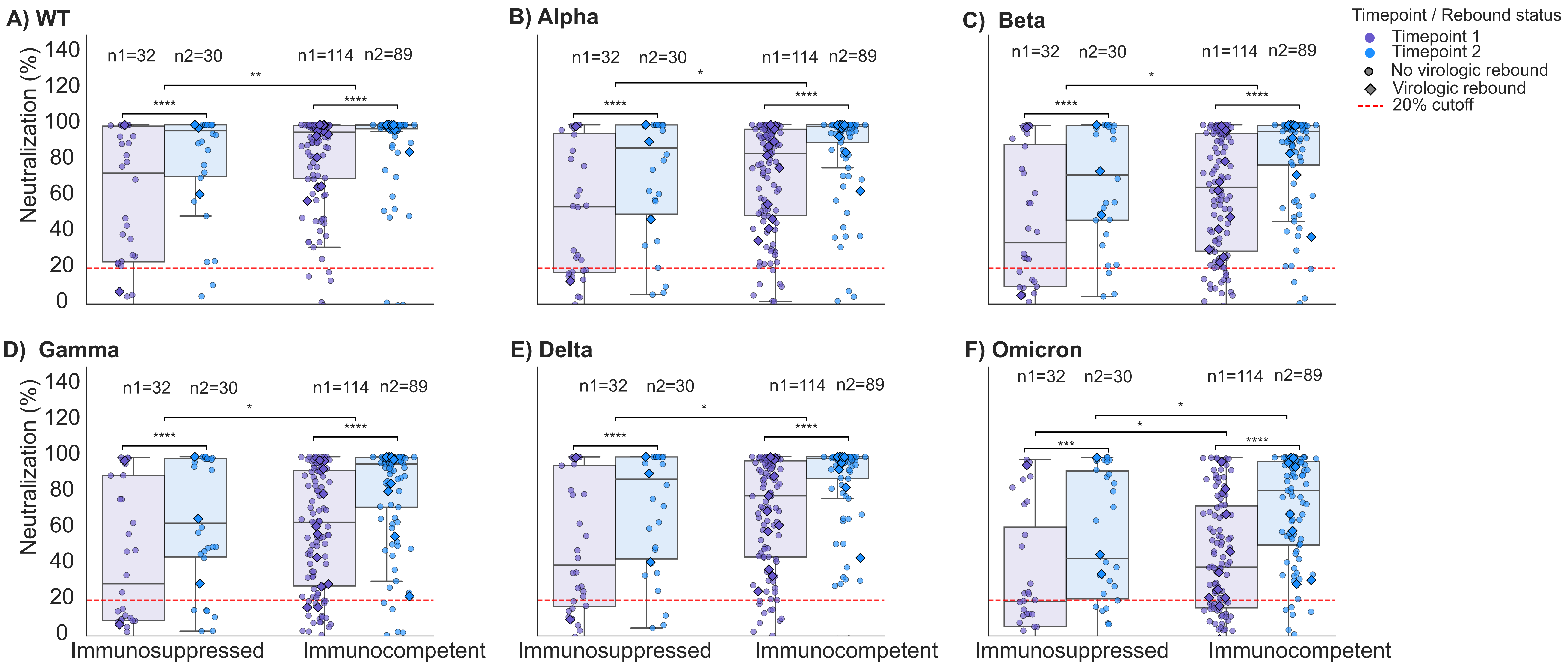


**Figure: Neutralizing antibody levels over time in participants who are immunocompromised and immunocompetent.** Serum neutralization of different variants of SARS-CoV-2 A) SARS-CoV-2 WT, B) SARS-CoV-2 B.1.1.7/ Alpha, C) SARS-CoV-2 B.1.351/Beta, D) SARS-CoV-2 P.1/Gamma, E) SARS-CoV-2 B.1.617.2/Delta and F) SARS-CoV-2 B.1.1.529/Omicron in Paxlovid treated and untreated participants in 2 different timepoints after the detection of COVID-19. Timepoint 1 represents 0-15 days after detection ( acute phase), Timepoint 2 represents 15 to 60 days after detection (post-acute phase). Sample sizes per timepoint (n1, n2 are indicated above each group. Multiple comparisons for different timepoints were performed using Wilcoxon matched-pairs signed rank test. Comparisons between independent groups were performed using the Mann-Whitney U test. Box plots show 25th, 50th, and 75th percentiles; whiskers, maximum and minimum. All data points are shown; y-axes show percentage. The diamond dots represent virologic rebounders. Symbols above the brackets indicate the degree of significance. No asterisks is nonsignificant, **** p<0.0001, *** P < 0.001, ** P < 0.01 and * P < 0.05.

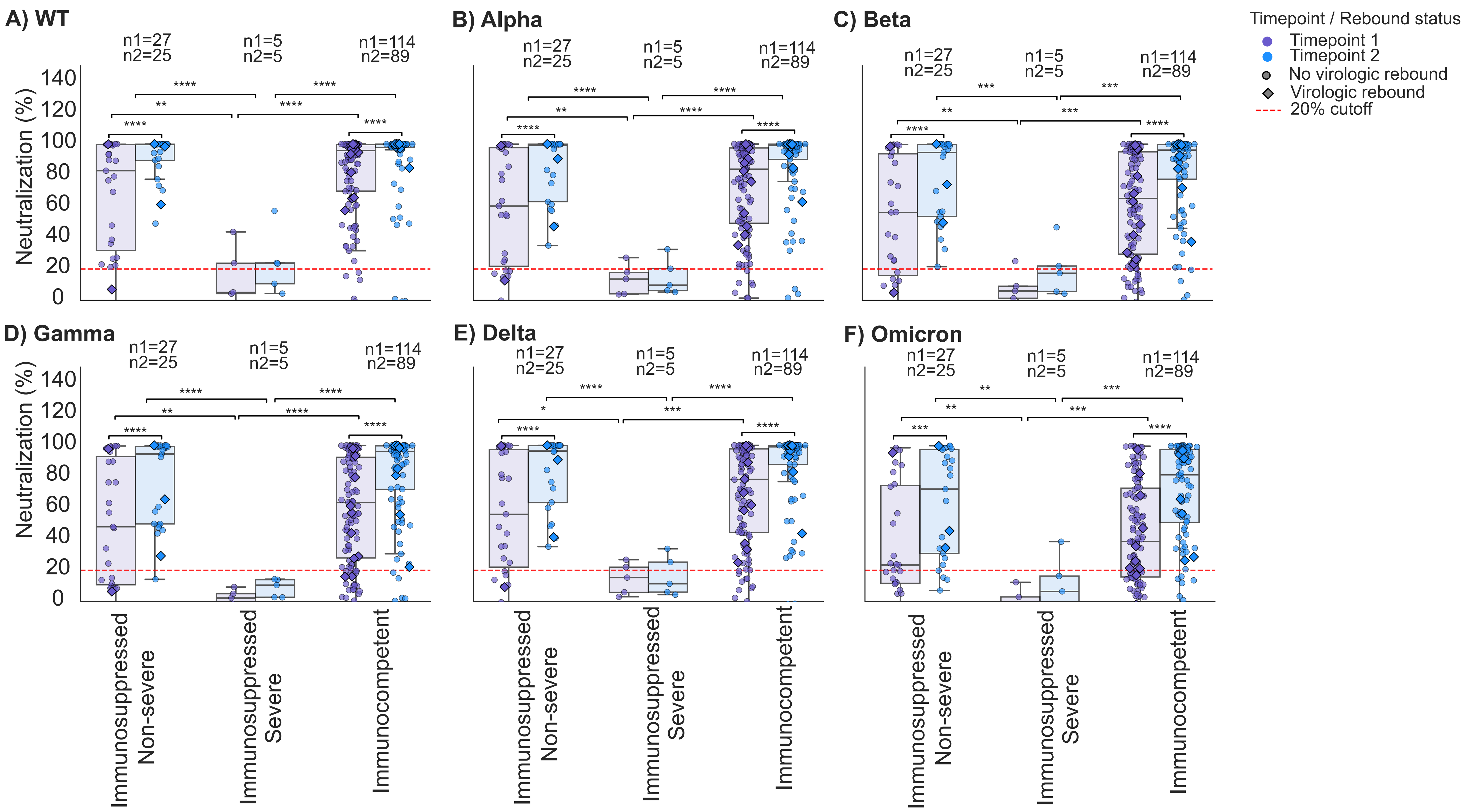


**Figure: Neutralizing antibody levels over time in participants with Non-severe immunosuppressed, severe immunosuppressed and immunocompetent.**
Serum neutralization of different variants of SARS-CoV-2 A) SARS-CoV-2 WT, B) SARS-CoV-2 B.1.1.7/ Alpha, C) SARS-CoV-2 B.1.351/Beta, D) SARS-CoV-2 P.1/Gamma, E) SARS-CoV-2 B.1.617.2/Delta and F) SARS-CoV-2 B.1.1.529/Omicron in the participants in 2 different timepoints after the detection of COVID-19. Timepoint 1 represents 0-15 days after detection ( acute phase), Timepoint 2 represents 15 to 60 days after detection (post-acute phase). Sample sizes per timepoint (n1, n2) are indicated above each group. Multiple comparisons for different timepoints were performed using Wilcoxon matched-pairs signed rank test. Comparisons between independent groups were performed using the Mann-Whitney U test. Box plots show 25th, 50th, and 75th percentiles; whiskers, maximum and minimum. All data points are shown; y-axes show percentage. The diamond dots represent virologic rebounders. Symbols above the brackets indicate the degree of significance. No asterisks is nonsignificant, **** p<0.0001, *** P < 0.001, ** P < 0.01 and * P < 0.05.

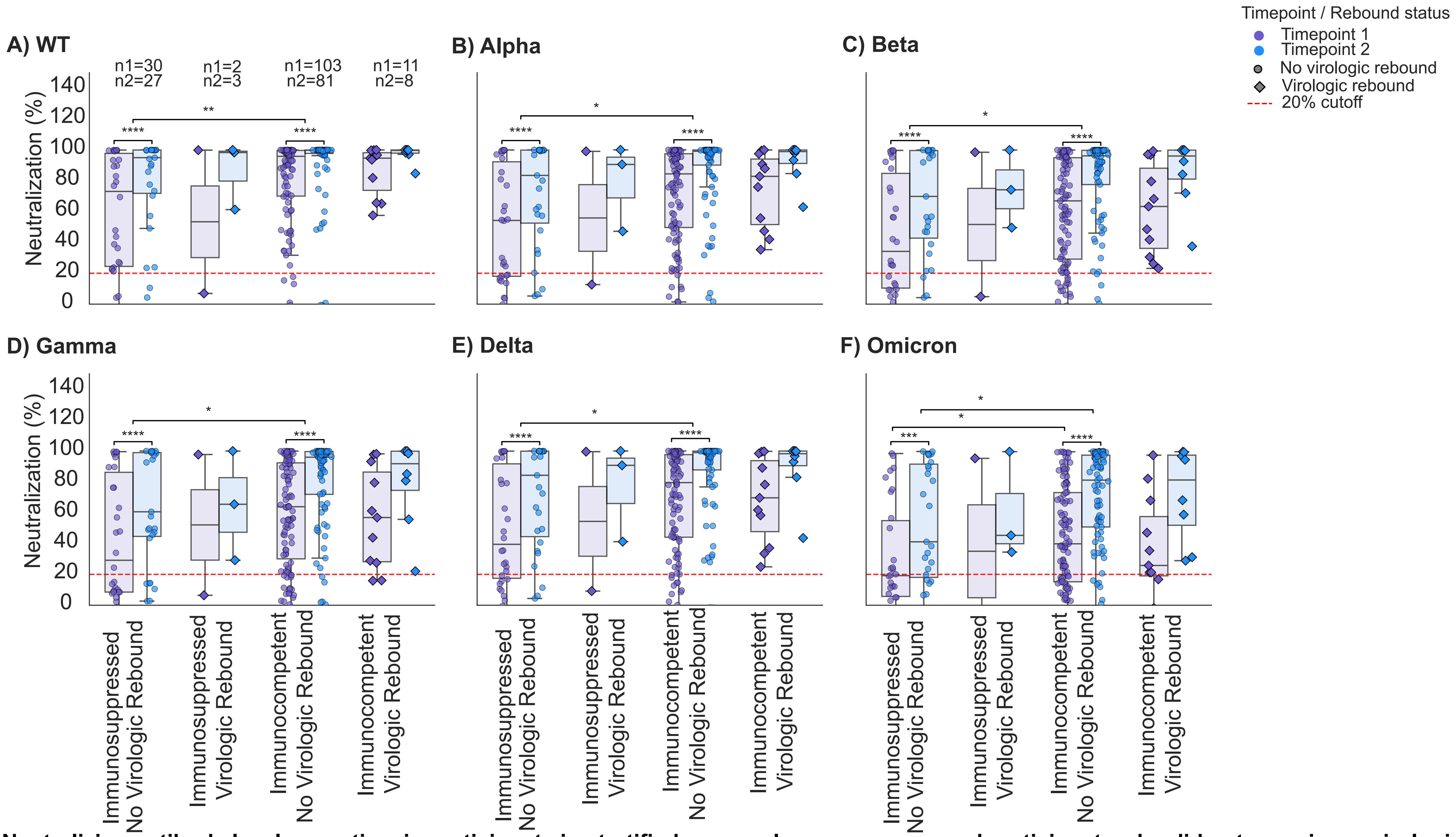


**Figure: Neutralizing antibody levels over time in participants in stratified groups: Immunosuppressed participants who did not experience virologic rebound, immunosuppressed participants who experienced virologic rebound, immunocompetant participants who did not experience virologic rebound and immunocompetant participants who experienced virologic rebound.** Serum neutralization ofdifferent variants of SARS-CoV-2 A) SARS-CoV-2 WT, B) SARS-CoV-2 B.1.1.7/ Alpha, C) SARS-CoV-2 B.1.351/Beta, D) SARS-CoV-2 P.1/Gamma, E) SARS-CoV-2 B.1.617.2/Delta and F) SARS-CoV-2 B.1.1.529/Omicron in The participants in 2 different timepoints after the detectionof COVID-19. Timepoint 1 represents 0-15 days after detection ( acute phase), Timepoint 2 represents 15 to 60 days after detection (post-acute phase). Sample sizes per timepoint (n1, n2 are indicated above each group. Multiple comparisons for different timepoints were performed using Wilcoxon matched-pairs signed rank test. Comparisons between independent groups were performed using the Mann-Whitney U test. Box plots show 25th, 50th, and 75th percentiles; whiskers, maximum and minimum. All data points are shown; y-axes show percentage. The diamond dots represent virologic rebounders. Symbols above the brackets indicate the degree of significance. No asterisks is nonsignificant, **** p<0.0001, *** P < 0.001, ** P < 0.01 and * P < 0.05.

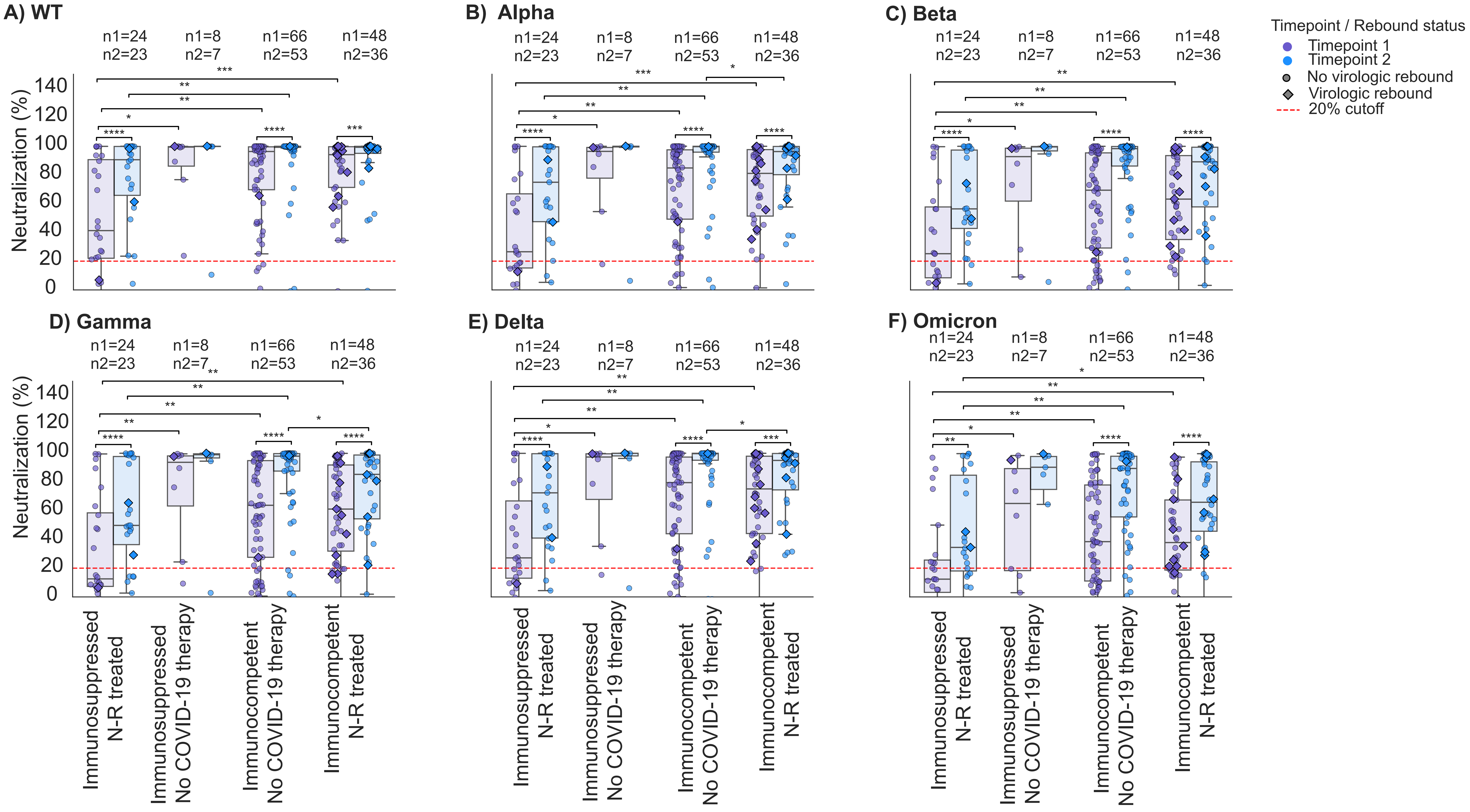


**Figure: Neutralizing antibody levels over time in participants in stratified groups: Immunosuppressed participants who received N-R treatment and who did not receive any therapy and immunocompetant participants who received N-R treatment and who did not receive any therapy.** Serum neutralization of different variants of SARS-CoV-2 A) SARS-CoV-2 WT, B) SARS-CoV-2 B.1.1.7/ Alpha, C) SARS-CoV-2 B.1.351/Beta, D) SARS-CoV-2 P.1/ Gamma, E) SARS-CoV-2 B.1.617.2/Delta and F) SARS-CoV-2 B.1.1.529/Omicron in the participants in 2 different timepoints after the detectionof COVID-19. Timepoint 1 represents 0-15 days after detection ( acute phase), Timepoint 2 represents 15 to 60 days after detection (post-acute phase). Sample sizes per timepoint (n1, n2 are indicated above each group. Multiple comparisons for different timepoints were performed using Wilcoxon matched-pairs signed rank test. Comparisons between independent groups were performed using the Mann-Whitney U test. Box plots show 25th, 50th, and 75th percentiles; whiskers, maximum and minimum. All data points are shown; y-axes show percentage. The diamond dots represent virologic rebounders. Symbols above the brackets indicate the degree of significance. No asterisks is nonsignificant, **** p<0.0001, *** P < 0.001, ** P < 0.01 and * P < 0.05.
