## Supplementary data part 2 for "Post-infection immune response in adults with COVID-19 with and without nirmatrelvir-ritonavir treatment and virologic rebound"

**Supplementaty Data Part 2**

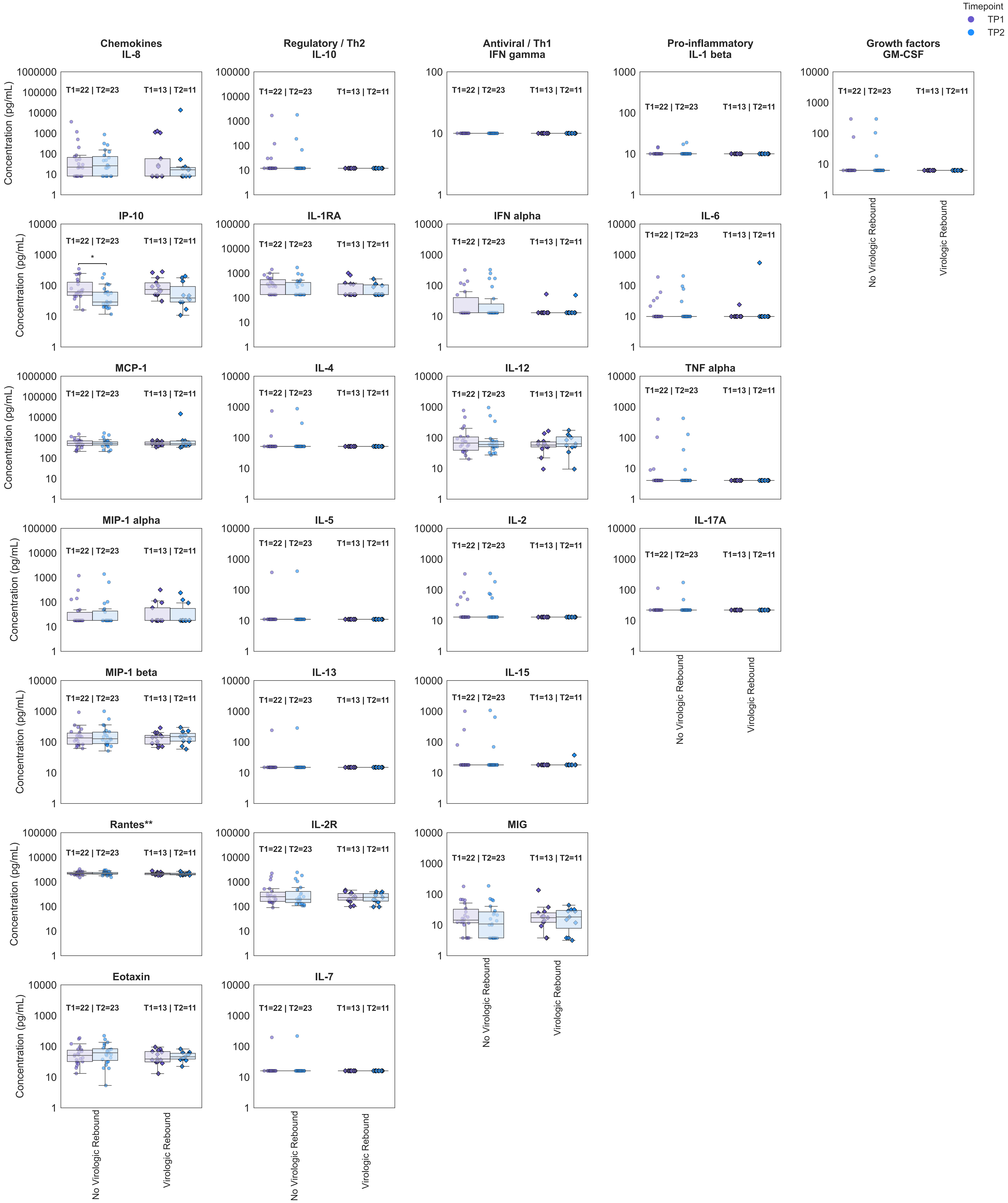


**Figure . Longitudinal serum cytokine and chemokine profiles stratified by Virologic Rebound status.**Cytokines were grouped according to biological function, including chemokines, regulatory/Th2 cytokines, antiviral/Th1 cytokines, pro-inflammatory cytokines, and growth factors. Individual data points represent participant samples collected at Timepoint 1 (TP1, purple) and Timepoint 2 (TP2, blue). Circles indicate participants without virologic rebound, whereas diamonds indicate participants with rebound. Boxplots display the median and interquartile range; whiskers represent 1.5× the interquartile range. Within-group longitudinal comparisons between TP1 and TP2 were performed using paired Wilcoxon signed-rank tests when paired samples were available. Between-group comparisons were performed using two-sided Mann–Whitney U tests.Statistical significance is indicated only for significant comparisons (*P < 0.05, **P < 0.01, ***P < 0.001, ****P < 0.0001). Sample sizes for each timepoint are indicated above each group.

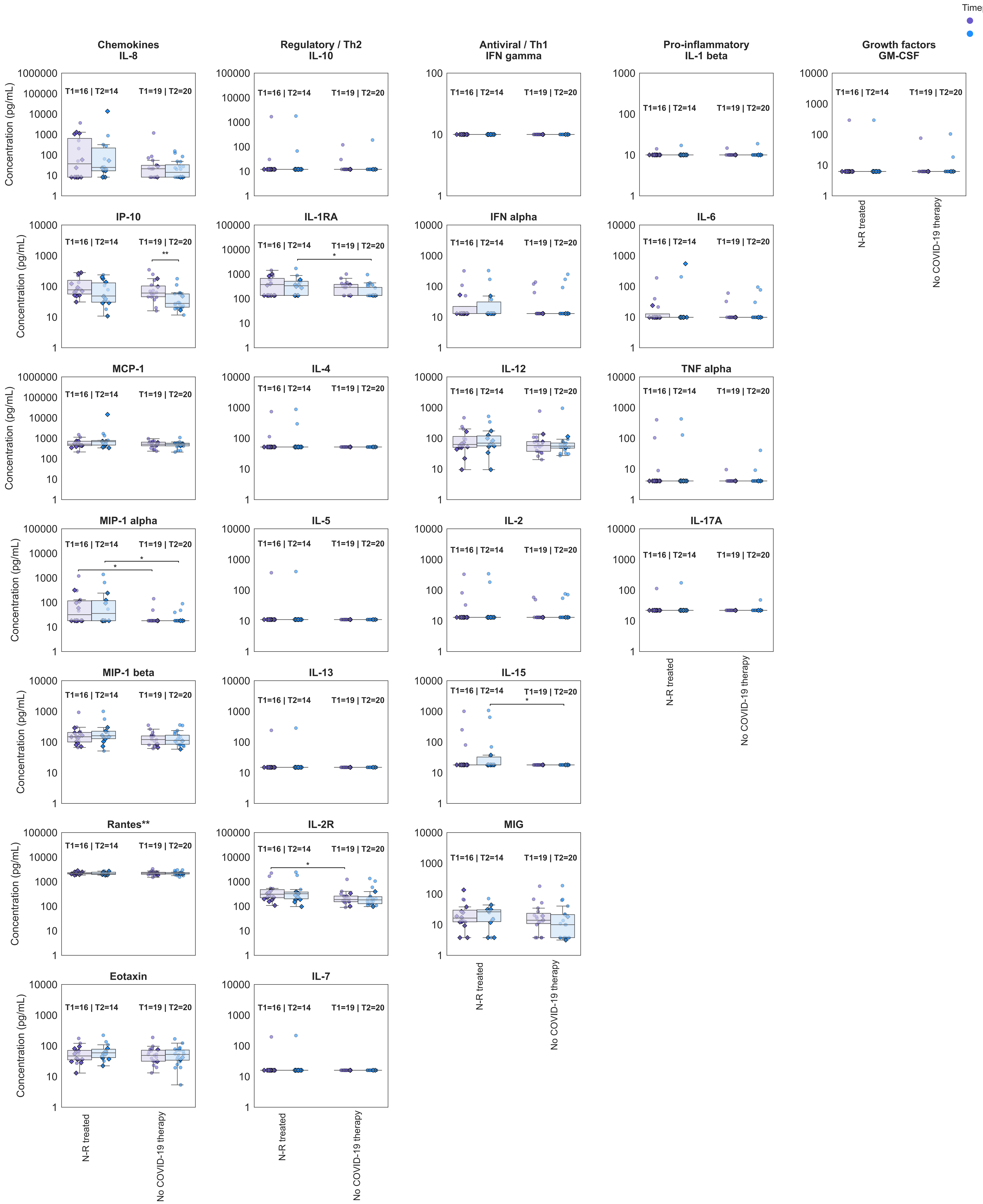


**Figure . Longitudinal serum cytokine and chemokine profiles stratified by COVID-19 treatment status.** Cytokines were grouped according to biological function, including chemokines, regulatory/Th2 cytokines, antiviral/Th1 cytokines, pro-inflammatory cytokines, and growth factors. Individual data points represent participant samples collected at Timepoint 1 (TP1, purple) and Timepoint 2 (TP2, blue). Circles indicate participants without virologic rebound, whereas diamonds indicate participants with rebound. Boxplots display the median and interquartile range; whiskers represent 1.5× the interquartile range. Within-group longitudinal comparisons between TP1 and TP2 were performed using paired Wilcoxon signed-rank tests when paired samples were available. Between-group comparisons were performed using two-sided Mann–Whitney U tests.Statistical significance is indicated only for significant comparisons (*P < 0.05, **P < 0.01, ***P < 0.001, ****P < 0.0001). Sample sizes for each timepoint are indicated above each group.

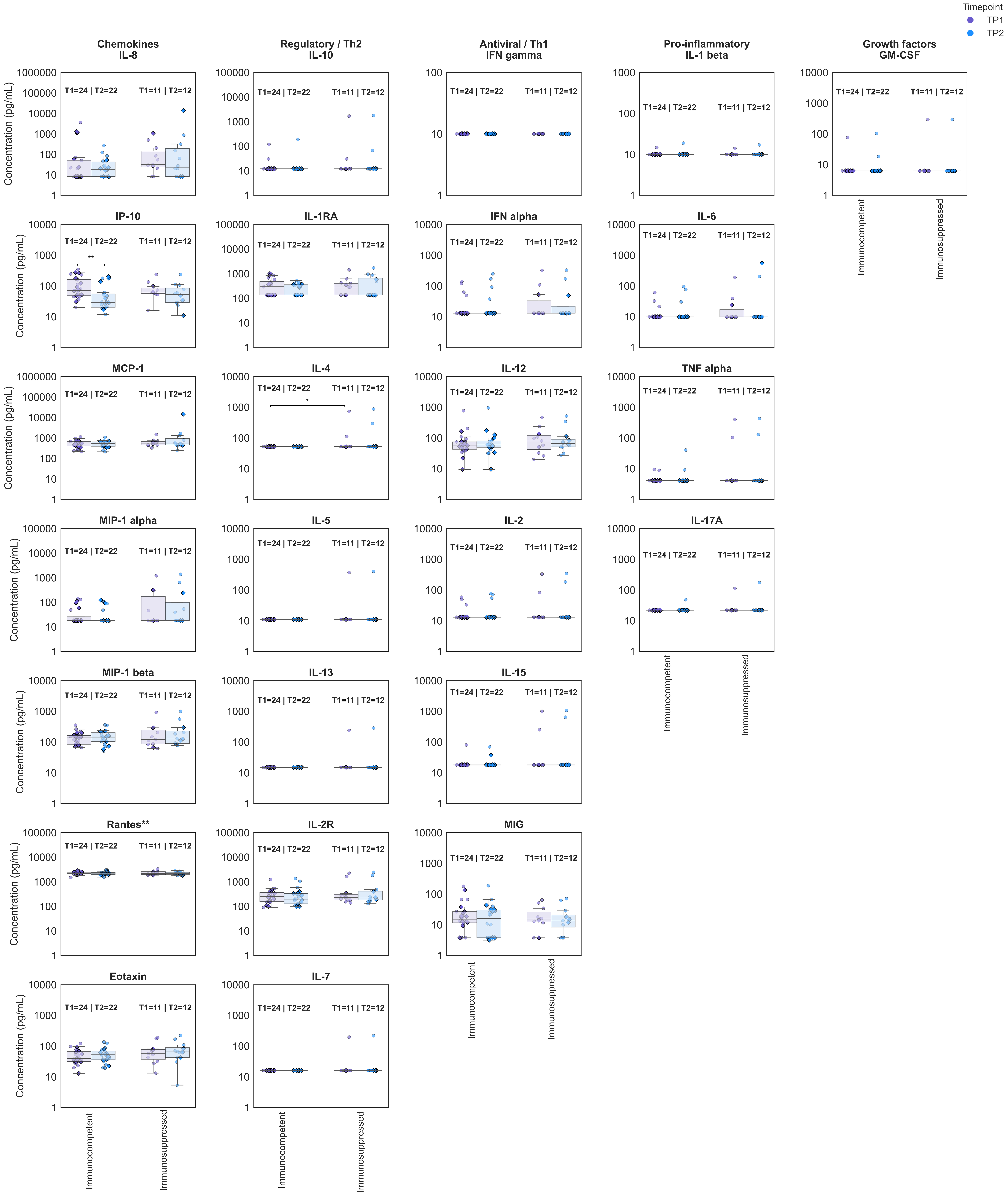


**Figure . Longitudinal serum cytokine and chemokine profiles stratified by Immune status.** Cytokines were grouped according to biological function, including chemokines, regulatory/Th2 cytokines, antiviral/Th1 cytokines, pro-inflammatory cytokines, and growth factors. Individual data points represent participant samples collected at Timepoint 1 (TP1, purple) and Timepoint 2 (TP2, blue). Circles indicate participants without virologic rebound, whereas diamonds indicate participants with rebound. Boxplots display the median and interquartile range; whiskers represent 1.5× the interquartile range. Within-group longitudinal comparisons between TP1 and TP2 were performed using paired Wilcoxon signed-rank tests when paired samples were available. Between-group comparisons were performed using two-sided Mann–Whitney U tests.Statistical significance is indicated only for significant comparisons (*P < 0.05, **P < 0.01, ***P < 0.001, ****P < 0.0001). Sample sizes for each timepoint are indicated above each group.

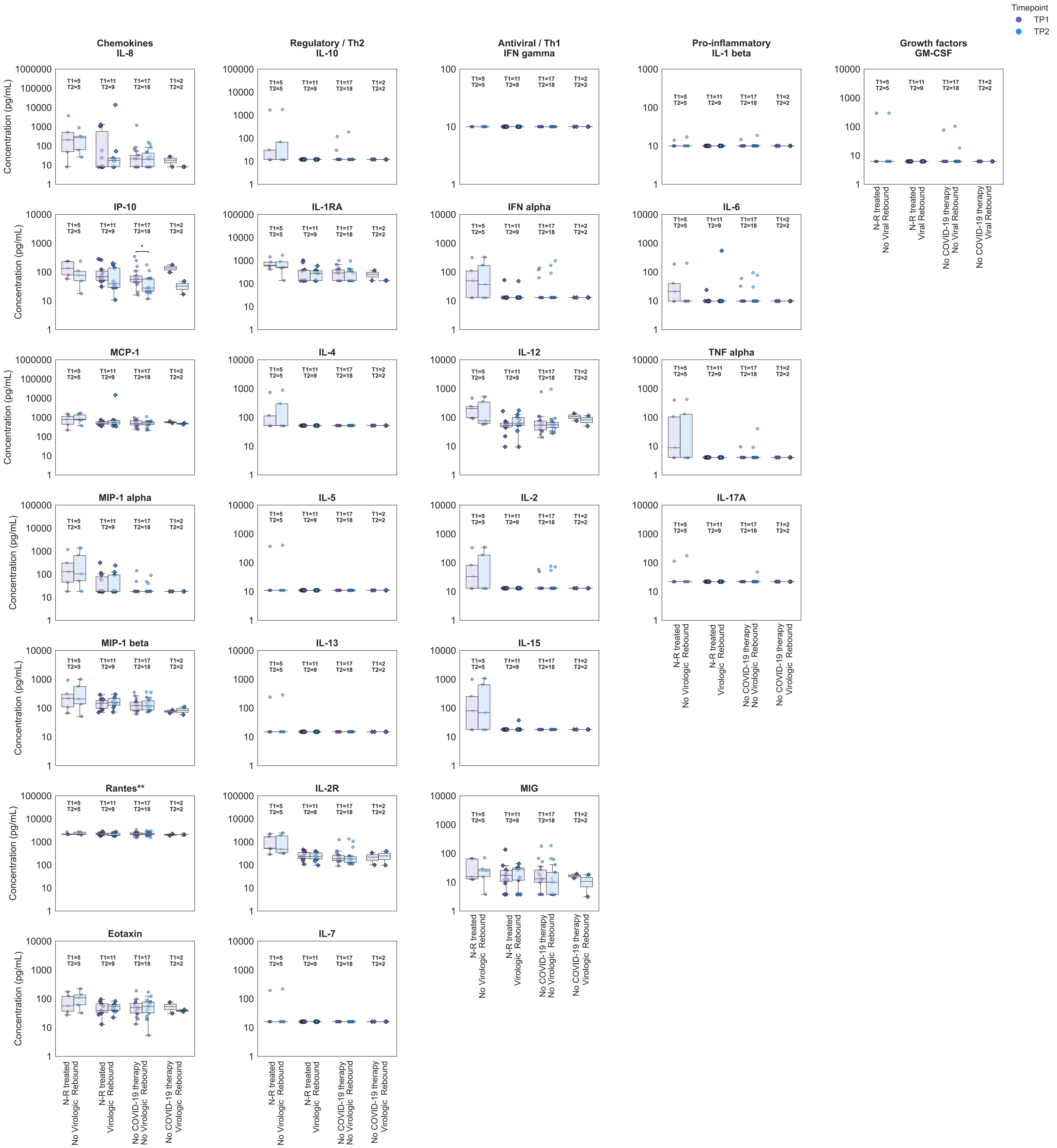


**Figure . Longitudinal serum cytokine and chemokine profiles stratified by Treatment and virologic rebound status.**Cytokines were grouped according to biological function, including chemokines, regulatory/Th2 cytokines, antiviral/Th1 cytokines, pro-inflammatory cytokines, and growth factors. Individual data points represent participant samples collected at Timepoint 1 (TP1, purple) and Timepoint 2 (TP2, blue). Circles indicate participants without virologic rebound, whereas diamonds indicate participants with rebound. Boxplots display the median and interquartile range; whiskers represent 1.5× the interquartile range. Within-group longitudinal comparisons between TP1 and TP2 were performed using paired Wilcoxon signed-rank tests when paired samples were available. Between-group comparisons were performed using two-sided Mann–Whitney U tests.Statistical significance is indicated only for significant comparisons (*P < 0.05, **P < 0.01, ***P < 0.001, ****P < 0.0001). Sample sizes for each timepoint are indicated above each group.

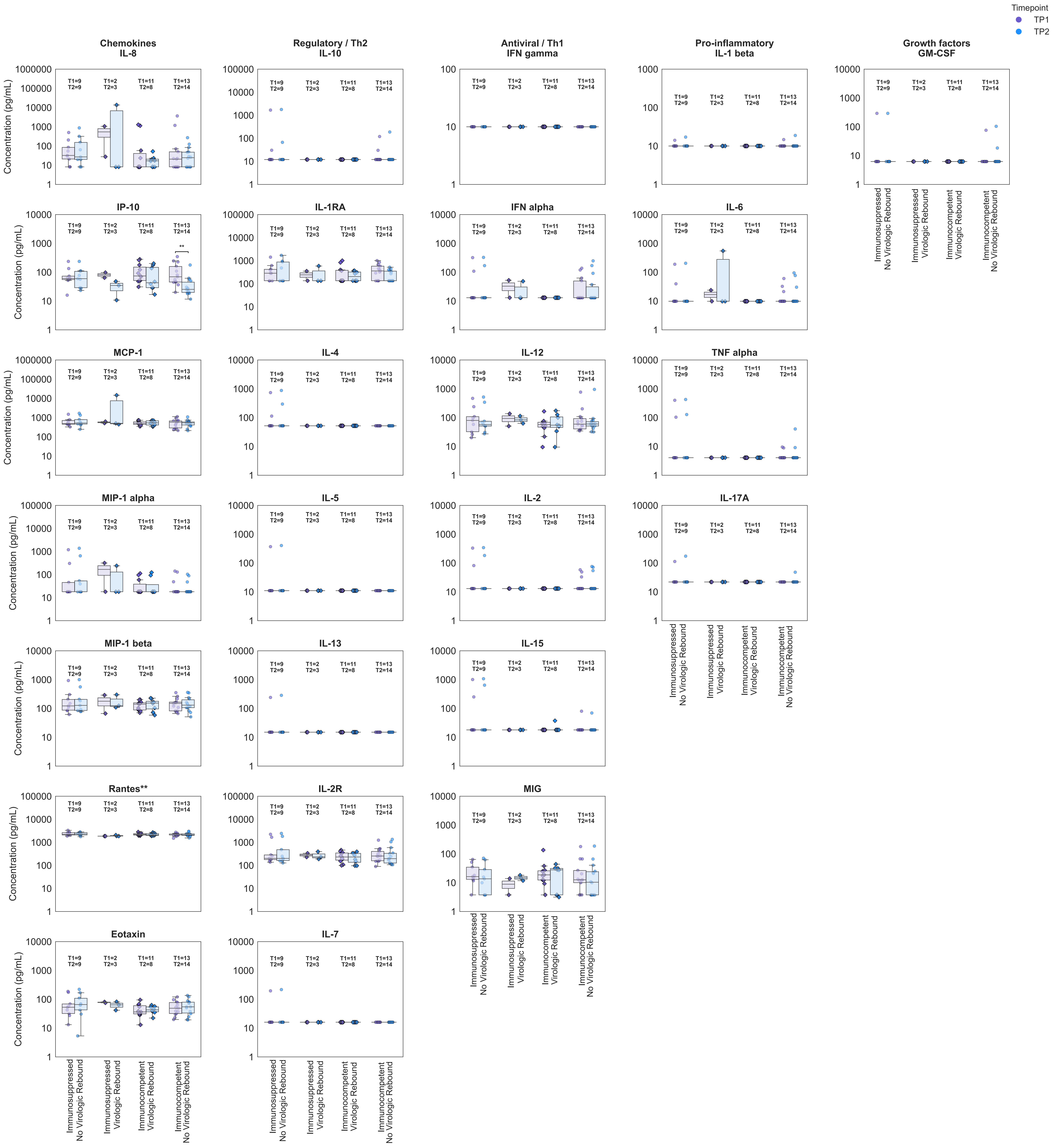


**Figure . Longitudinal serum cytokine and chemokine profiles stratified by Immune and virologic rebound status.**Cytokines were grouped according to biological function, including chemokines, regulatory/Th2 cytokines, antiviral/Th1 cytokines, pro-inflammatory cytokines, and growth factors. Individual data points represent participant samples collected at Timepoint 1 (TP1, purple) and Timepoint 2 (TP2, blue). Circles indicate participants without virologic rebound, whereas diamonds indicate participants with rebound. Boxplots display the median and interquartile range; whiskers represent 1.5× the interquartile range. Within-group longitudinal comparisons between TP1 and TP2 were performed using paired Wilcoxon signed-rank tests when paired samples were available. Between-group comparisons were performed using two-sided Mann–Whitney U tests.Statistical significance is indicated only for significant comparisons (*P < 0.05, **P < 0.01, ***P < 0.001, ****P < 0.0001). Sample sizes for each timepoint are indicated above each group.

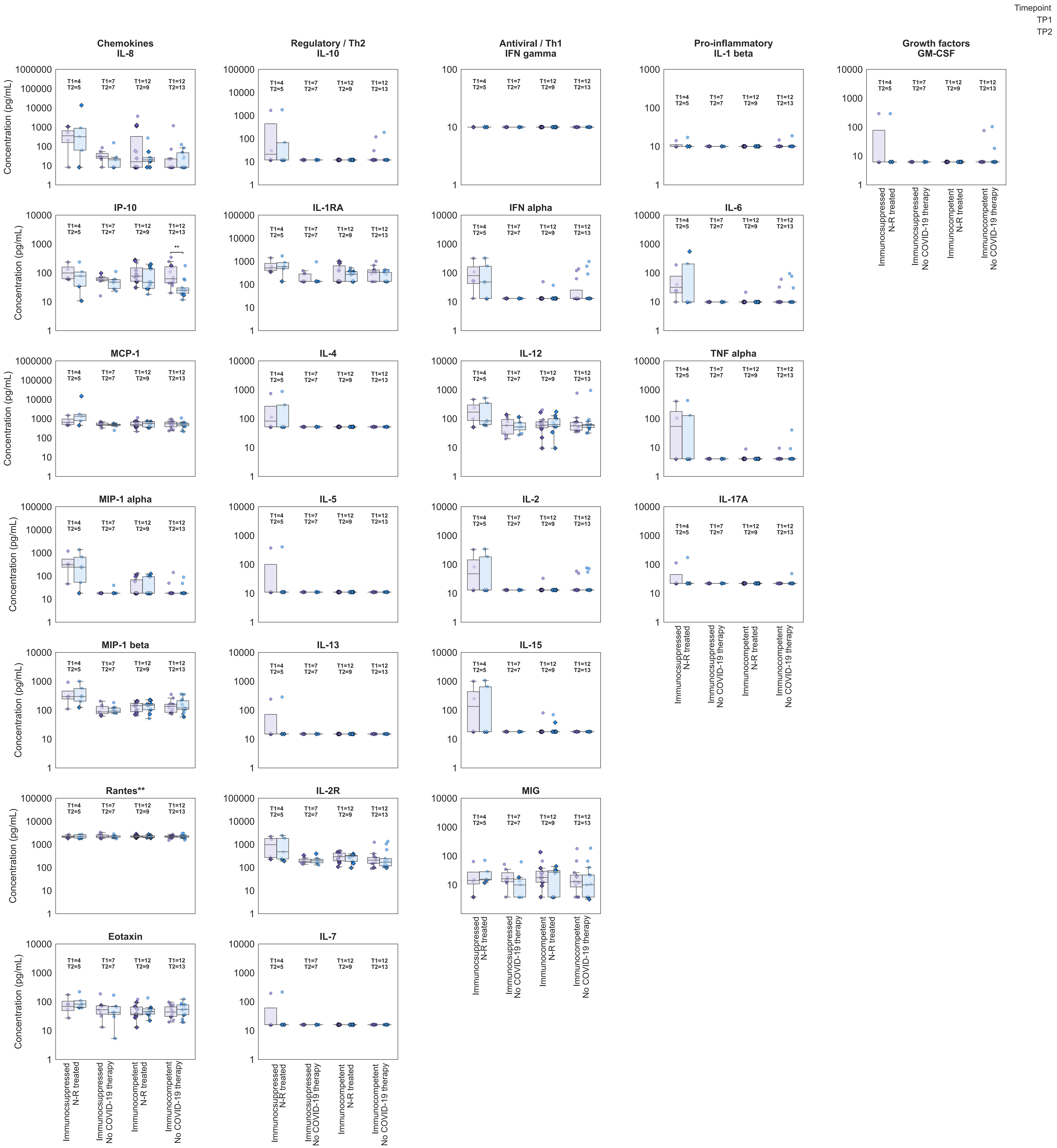


**Figure . Longitudinal serum cytokine and chemokine profiles stratified by Immune and Treatment status.**Cytokines were grouped according to biological function, including chemokines, regulatory/Th2 cytokines, antiviral/Th1 cytokines, pro-inflammatory cytokines, and growth factors. Individual data points represent participant samples collected at Timepoint 1 (TP1, purple) and Timepoint 2 (TP2, blue). Circles indicate participants without virologic rebound, whereas diamonds indicate participants with rebound. Boxplots display the median and interquartile range; whiskers represent 1.5× the interquartile range. Within-group longitudinal comparisons between TP1 and TP2 were performed using paired Wilcoxon signed-rank tests when paired samples were available. Between-group comparisons were performed using two-sided Mann–Whitney U tests.Statistical significance is indicated only for significant comparisons (*P<0.05, **P<0.01,***P<0.001, ****P< 0.0001). Sample sizes for each timepoint are indicated above each group.
